# Multiscale activity imaging in mammary gland reveals how oxytocin enables lactation

**DOI:** 10.1101/657510

**Authors:** Alexander J. Stevenson, Gilles Vanwalleghem, Teneale A. Stewart, Nicholas D. Condon, Bethan Lloyd-Lewis, Natascia Marino, James W. Putney, Ethan K. Scott, Adam D. Ewing, Felicity M. Davis

## Abstract

The mammary epithelium is indispensable for the continued survival of more than 5000 mammalian species. For some, the volume of milk ejected in a single day exceeds their entire blood volume. Here, we unveil the spatiotemporal properties of physiological signals that orchestrate milk ejection. Using quantitative, multidimensional imaging of mammary cell ensembles, we reveal how stimulus evoked Ca^2+^ oscillations couple to contractions in basal epithelial cells. Moreover, we show that Ca^2+^-dependent contractions generate the requisite force to physically deform the innermost layer of luminal cells, compelling them to discharge the fluid that they produced and housed. Through the collective action of thousands of these biological positive displacement pumps, each linked to a contractile ductal network, milk is delivered into the mouth of the dependent neonate, seconds after the command.

**Significance Statement:** The mammary gland is functional for only a brief period of a female’s lifetime—if at all. During this time, it operates not for the survival of the individual, but for the survival of her species. Here, we visualize the nature of alveolar contractions in the functionally-mature mammary gland, revealing how specialized epithelial cells, which possess the ability to behave like smooth muscle cells, undergo Ca^2+^-dependent contractions. We demonstrate that individual oscillators can be electrically coupled to achieve global synchrony, a phenomenon that has not yet been observed in the mammary gland. By imaging activity across scales, we provide a window into the organization, dynamics and role of epithelial Ca^2+^ oscillations in the organ principally responsible for sustaining neonatal life in mammals.

## Introduction

The ability to visualize how a single living cell, in its native environment, translates an extracellular message into an intracellular signal to execute a defined task at the cell level and cooperatively achieve a biological outcome at the organ level is revolutionizing our understanding of multicellular systems. Such an approach has provided new insights into a range of biological phenomena, including how plants defend against herbivory (1), how fish escape looming predators (2, 3) and how mammals store memories (4). The rational design and continued refinement of genetically-encoded Ca^2+^ indicators (GECIs) has fueled these advances (5). However, the use of GECIs for *in situ* activity mapping in adult vertebrates, has largely remained an achievement of neuroscience, where neural activity is tightly coupled to intracellular Ca^2+^ ([Ca^2+^]_i_) signaling (6). Efforts to map activity networks in specific populations of non-excitable cells in other solid organs is lagging. Our understanding of how epithelial tissues function, for example, has principally arisen through analysis of isolated cells (often serially propagated under physiologically extraneous conditions), retrospective examination of fixed tissue and interrogation of genetic knockout models (where biological function is inferred in the absence of physiological redundancy or compensation).

The mammary gland has a universal and indispensable role in mammalian offspring survival. In its functionally-mature state, it consists of an inner layer of luminal (milk producing) epithelial cells and an outer layer of contractile basal epithelial cells (also known as myoepithelial cells) (7). When young offspring suckle, maternally produced oxytocin (OT) binds to its cognate receptor (the OXTR, a Gq-linked G-protein coupled receptor) on mammary basal cells, causing them to contract (8). A model, therefore, emerges where activity may be tightly coupled to [Ca^2+^]_i_ in this organ—via phospholipase C (PLC)-inositol trisphosphate (InsP3) signaling (8–12)—making functional *in situ* imaging of an epithelial signal-response relationship possible.

Here, we engineered mice with directed expression of a GECI to basal epithelial cells in the mammary gland. This enabled us to quantitatively probe the organization and function of real-time [Ca^2+^]_i_ signaling events in individual cells within this complex living tissue, at a level of rigor that has only previously been achieved in the adult brain.

## Results

### Basal cell [Ca^2+^]_i_ oscillations signal to repetitively deform mammary alveoli and force milk out

We developed transgenic mice that express the fast, ultrasensitive GECI GCaMP6f (5) under the inducible control of the cytokeratin (K) 5 gene promoter (13) (*GCaMP6f;K5CreERT2* mice) (Fig. 1A). The relatively high baseline fluorescence of this GECI is well suited for the quantitative assessment of [Ca^2+^]_i_ responses in functionally-mature basal cells, which are sparsely distributed with thin cellular processes (5, 14) (Fig. S1). GCaMP6f consists of a circularly permuted green fluorescent protein (GFP), enabling 3D assessment of its expression and lineage specific localization using an anti-GFP antibody (15) and optimized methods for tissue clearing (16). Genetic recombination in this model was high (Fig. S2) and showed lineage restriction to basal epithelial cells (Fig. 1B).

**Figure 1.**
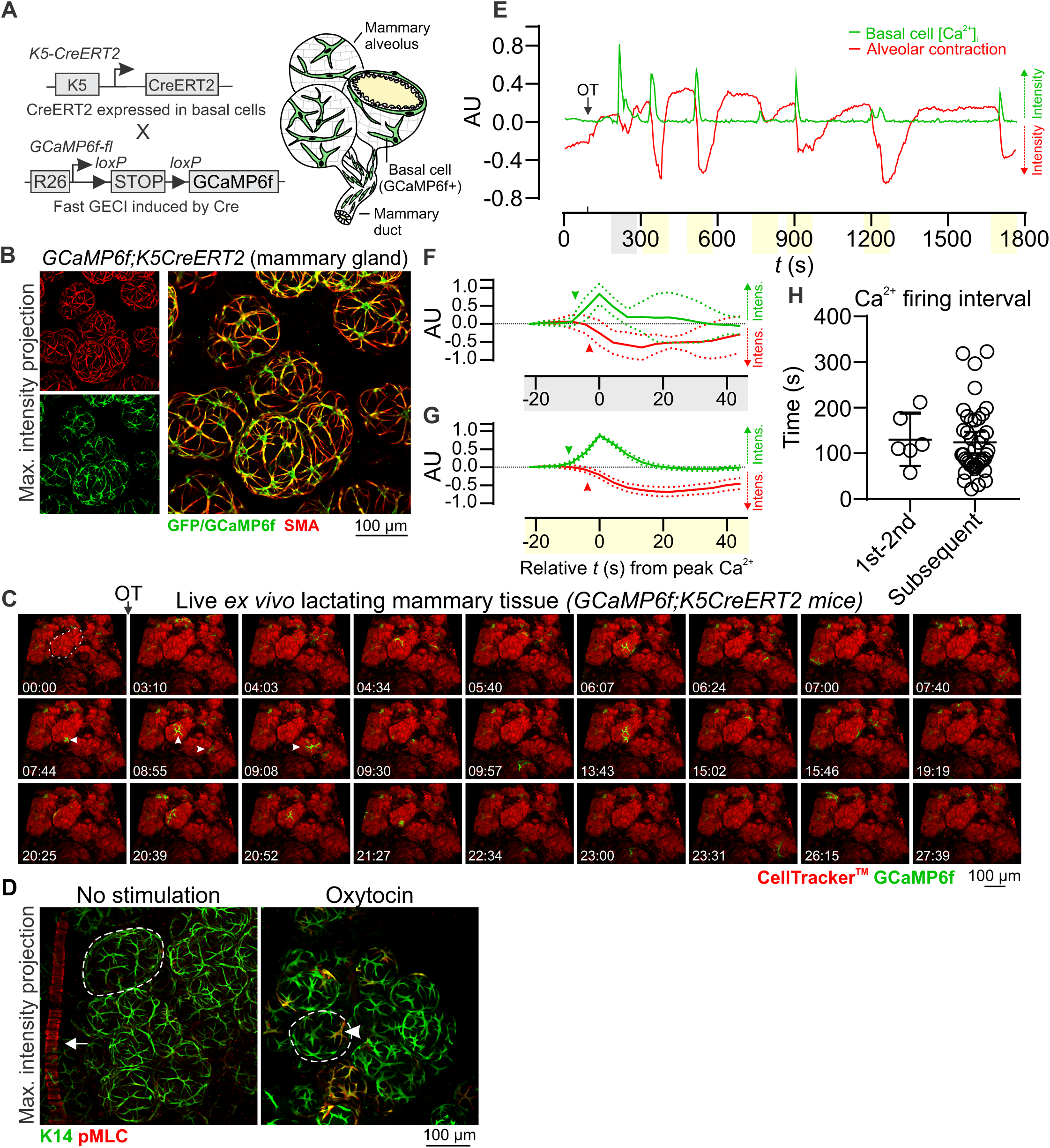
Basal cell Ca^2+^ oscillations precede alveolar contractions. (**A**) Schematic representation of the *GCaMP6f;K5CreERT2* model. (**B**) Maximum intensity z-projection (0-60 μm) of cleared lactating mammary tissue immunostained with smooth muscle actin (SMA) to reveal basal cells and an anti-GFP antibody to detect GCaMP6f expression. (**C**) 3D time-lapse imaging of live mammary tissue from *GCaMP6f;K5CreERT2* lactating mice stimulated with OT (85 nM) at 01:33 (min:s). Images show maximum intensity z-projection 32 μm through live tissue. Dotted line (frame 1) identifies a single alveolar unit; arrowheads point to example Ca^2+^ events in single cells. See also Movie S1. (**D**) Maximum intensity z-projections [0-120 μm (left), 0-44 μm (right)] of cleared mammary tissue immunostained with K14 to reveal basal cells and pMLC to show sites of contractile activity. Arrow shows pMLC^+^ blood vessel in control mammary tissue (lactation), arrowhead shows pMLC^+^ basal cell in lactating mammary tissue stimulated with OT (85 nM) prior to fixation; dotted lines surround single alveolar units. (**E**) Quantification of [Ca^2+^]_i_ responses (green) and alveolar unit contraction (red) in lactating mammary tissue from *GCaMP6f;K5CreERT2* mice. [Ca^2+^]_i_ measurements are ΔF/F0. Alveolar unit contractions are shown by negative deflections (CellTracker™ red fluorescence). (**F-G**) Average (± SEM) peak [Ca^2+^]_i_ and contractile responses. Highlighting (*x*-axis) corresponds with events linked in E; arrowheads show initiation of the response. (**H**) Interval between the first-second and all subsequent [Ca^2+^]_i_ events. AU, arbitrary unit; n = 3 mice.

To assess OT-mediated basal cell [Ca^2+^]_i_ responses, we performed 4-dimensional (*x*-, *y*-, *z*-, *t*-) quantitative imaging of *ex vivo* mammary tissue pieces from lactating *GCaMP6f;K5CreERT2* mice—a method similar to the preparation of acute brain slices for neural imaging (17). Tissue was loaded with the live cell permeable dye CellTracker™ Red to visualize alveolar luminal (milk producing) cells. A large coordinated wave of [Ca^2+^]_i_, due to InsP3-mediated endoplasmic reticulum (ER) Ca^2+^ store release (8, 9, 12), was observed in mammary basal cells following OT stimulation and its diffusion through the tissue (Fig. 1C and **Movie S1**). This initial transient [Ca^2+^]_i_ elevation was followed by a phase of stochastic [Ca^2+^]_i_ oscillations (Fig. 1C and **Movie S1**) that were likely to be sustained in-part by Ca^2+^ influx across the plasma membrane (9, 11, 18).

The organization of basal cell contractions was also examined using 3-dimensional, deep tissue imaging of myosin light chain (MLC) phosphorylation. In tissue treated with OT prior to fixation, phospho-MLC (pMLC) -positive and -negative basal cells were observed to be interspersed throughout alveolar clusters (Fig. 1D), supporting the ostensibly stochastic nature of the mammary contractile response. Regions containing clusters of pMLC-positive cells, however, were occasionally observed (Fig. S3). Intravital imaging of OT-mediated [Ca^2+^]_i_ responses (19), supported observations in acute *ex vivo* tissue preparations (Fig S4 and **Movie S2**).

To determine whether increases in [Ca^2+^]_i_ are temporally correlated with alveolar unit contractions we quantified Ca^2+^-contraction responses in alveolar tissue. Whilst cell- and tissue-level movement is physiologically relevant and important, it poses additional computational challenges to the analysis of single cell Ca^2+^ responses in 4D image sequences. To overcome this, we utilized the diffeomorphic registration approach of Advanced Normalization Tools for motion correction (20, 21) (see Methods). This approach corrected major tissue movements, however, alveolar unit contractions remained largely intact, enabling quantification of [Ca^2+^]_i_ responses in basal cells and analysis of the physical distortions to the alveolar units that these cells embrace. These analyses confirmed that increases in [Ca^2+^]_i_ in individual basal cells were temporally correlated with physical distortions to the mechanically compliant luminal cell layer (Fig. 1E and Fig. S5). For both the first InsP3 response (Fig. 1F) and the subsequent oscillatory phase (Fig. 1G), increases in [Ca^2+^]_i_ preceded alveolar unit contractions. No statistical difference in the firing interval for [Ca^2+^]_i_ was observed between the first and second events and all subsequent events (Fig. 1H). These results reveal that each mammary alveolar unit, acting downstream of a basal cell OT-OXTR-InsP3-Ca^2+^ signaling axis, serves as a biological positive-displacement pump, repeatedly forcing milk out of its central lumen for passage through the ductal network.

### Basal cell contractions are Ca^2+^ signal dependent

To directly assess Ca^2+^-contraction coupling in mammary basal cells, we engineered triple transgenic mice that express GCaMP6f and the red fluorescent protein TdTomato (22) in basal cells (*GCaMP6f-TdTom;K5CreERT2* mice) (Fig. 2A). Using this model, we observed increases in [Ca^2+^]_i_ in single TdTomato-positive basal cells in response to OT, which immediately preceded their contraction (Fig. 2A-C and **Movie S3**). These data reveal with greater optical clarity how basal cells contract to deform the inner luminal cell layer for milk ejection and show unequivocally a temporal relationship between the Ca^2+^ signal and the contractile response.

**Figure 2.**
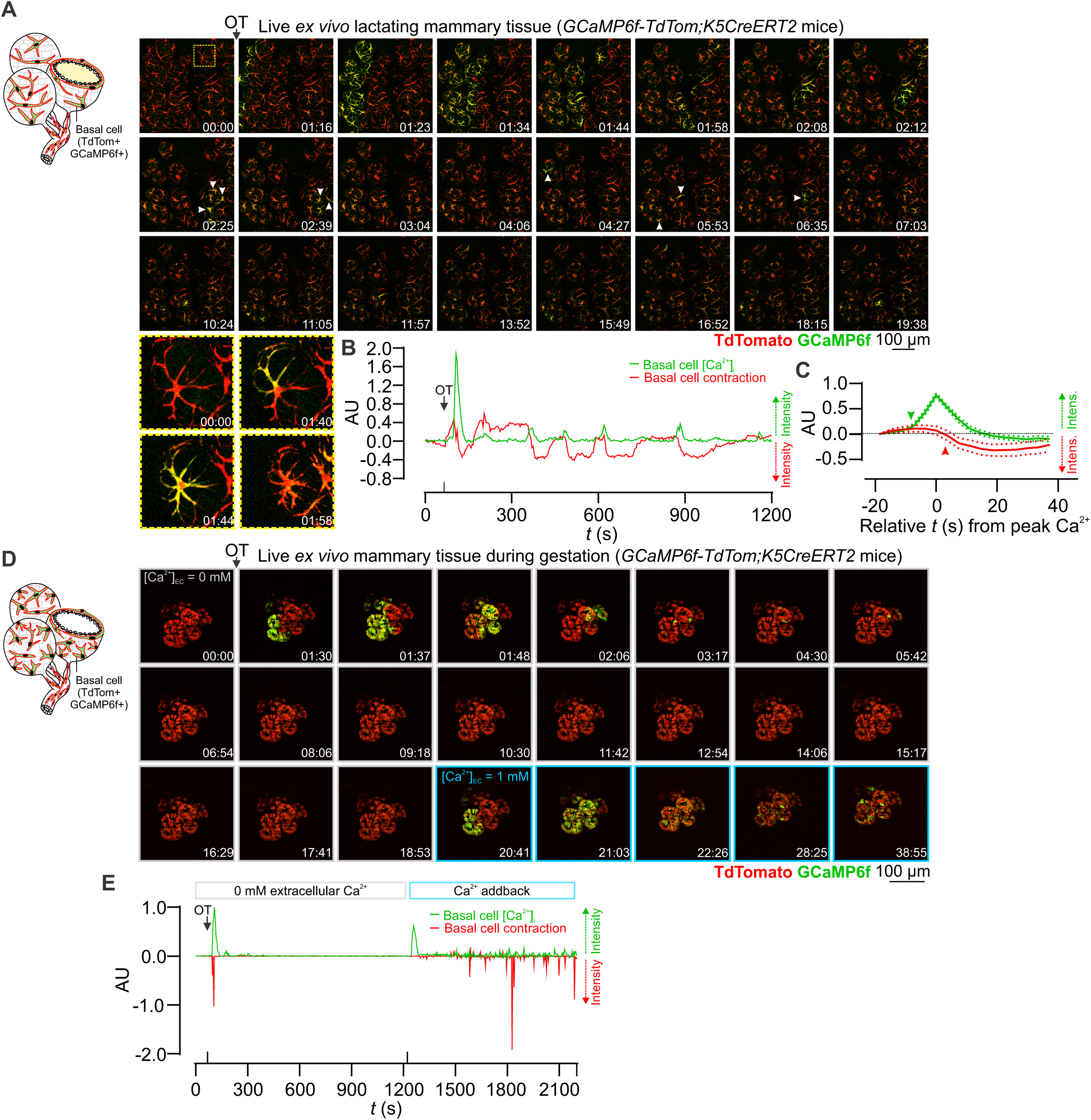
Ca^2+^-contraction coupling. (**A**) 3D time-lapse imaging of live mammary tissue from *GCaMP6f-TdTom;K5CreERT2* mice stimulated with OT (85 nM) at 01:09 (min:s). Images show maximum intensity z-projection (31.5 μm). Box (frame 1) expanded in panel below; arrowheads point to example Ca^2+^ events in single cells. See also **Movie S3**. (**B**) Quantification of [Ca^2+^]_i_ responses (green) and alveolar unit contraction (red) in lactating mammary tissue from *GCaMP6f-TdTom;K5CreERT2* mice. [Ca^2+^]_i_ measurements are ΔF/F0. Basal cell contractions are shown by negative deflections (TdTomato fluorescence). (**C**) Average (± SEM) peak [Ca^2+^]_i_ response (green) and contractile response (red) in mammary tissue isolated from lactating *GCaMP6f-TdTom;K5CreERT2* mice. Values were averaged from both the first response and the oscillatory phase. (**D**) 3D time-lapse imaging of live mammary tissue from *GCaMP6f-TdTom;K5CreERT2* mice (15.5-16.5 d.p.c., days post coitus) stimulated with OT (85 nM) at 01:08 (min:s) under extracellular Ca^2+^ free conditions. Images show maximum intensity z-projection (36 μm). Ca^2+^ (1 mM free) was added back at 20:23 (min:sec); extended recording conditions. See also **Movie S5**. (**E**) Quantification of [Ca^2+^]_i_ responses (green) and alveolar unit contraction (red) in mammary tissue from pregnant *GCaMP6f-TdTom;K5CreERT2* mice stimulated with OT under extracellular Ca^2+^ free conditions and with Ca^2+^ addback. [Ca^2+^]_i_ measurements are ΔF/F0. Basal cell contractions are shown by negative deflections (Jacobian determinant of the warp matrix for the TdTomato fluorescence). AU, arbitrary unit; n = 3 mice.

To determine whether Ca^2+^ forms an essential component of the signal transduction pathway linking OXTR engagement to basal cell contraction, we examined Ca^2+^-contraction responses under extracellular Ca^2+^-free conditions. Tissue was isolated from pregnant *GCaMP6fTdTom;K5CreERT2* mice and incubated in Ca^2+^-free physiological salt solution supplemented with the Ca^2+^ chelator BAPTA. By performing experiments using mammary tissue harvested prior to secretory activation (gestation day 15.5-16.5), when Ca^2+^-contraction coupling is observed (**Movie S4**), we were able to avoid the exceedingly high (> 90 mM) extracellular Ca^2+^ concentrations present in secreted milk (9). Under these conditions, addition of OT resulted in intracellular Ca^2+^ store release associated with cell contraction (Fig. 2D-E and **Movie S5**). Ensuing spike trains, however, were absent and subsequent contractions were abolished. Re-addition of extracellular Ca^2+^ led to the resumption of Ca^2+^ firing and basal cell contractions (Fig. 2D-E). These data demonstrate that both Ca^2+^ release from InsP3-sensitive (12) intracellular Ca^2+^ stores and Ca^2+^ influx across the plasma membrane are sufficient for basal cell contraction but that influx across the membrane is necessary to sustain cell and tissue contractions.

### Both ducts and alveoli contract to expel milk in the mature gland

The lactating mouse mammary gland consists of milk producing alveoli that are connected to the nipple via a branching ductal network (Fig. 1A). Heterogeneity in the expression of contractile markers in basal cells of ducts and alveoli has led to speculation that these two related (but spatially- and morphologically-distinct) cell populations are functionally divergent (23). We compared expression of myosin light chain kinase (MLCK), calponin (CNN1) and caldesmon (CALD1)—key components of the vascular smooth muscle contraction pathway that are upregulated in the mammary gland during lactation (Fig. S6)—in ducts versus alveoli of lactating mice (Fig. 3A) and humans (Fig. 3B). Our data reveal that these proteins are expressed at comparable levels in basal cells of both structures.

**Figure 3.**
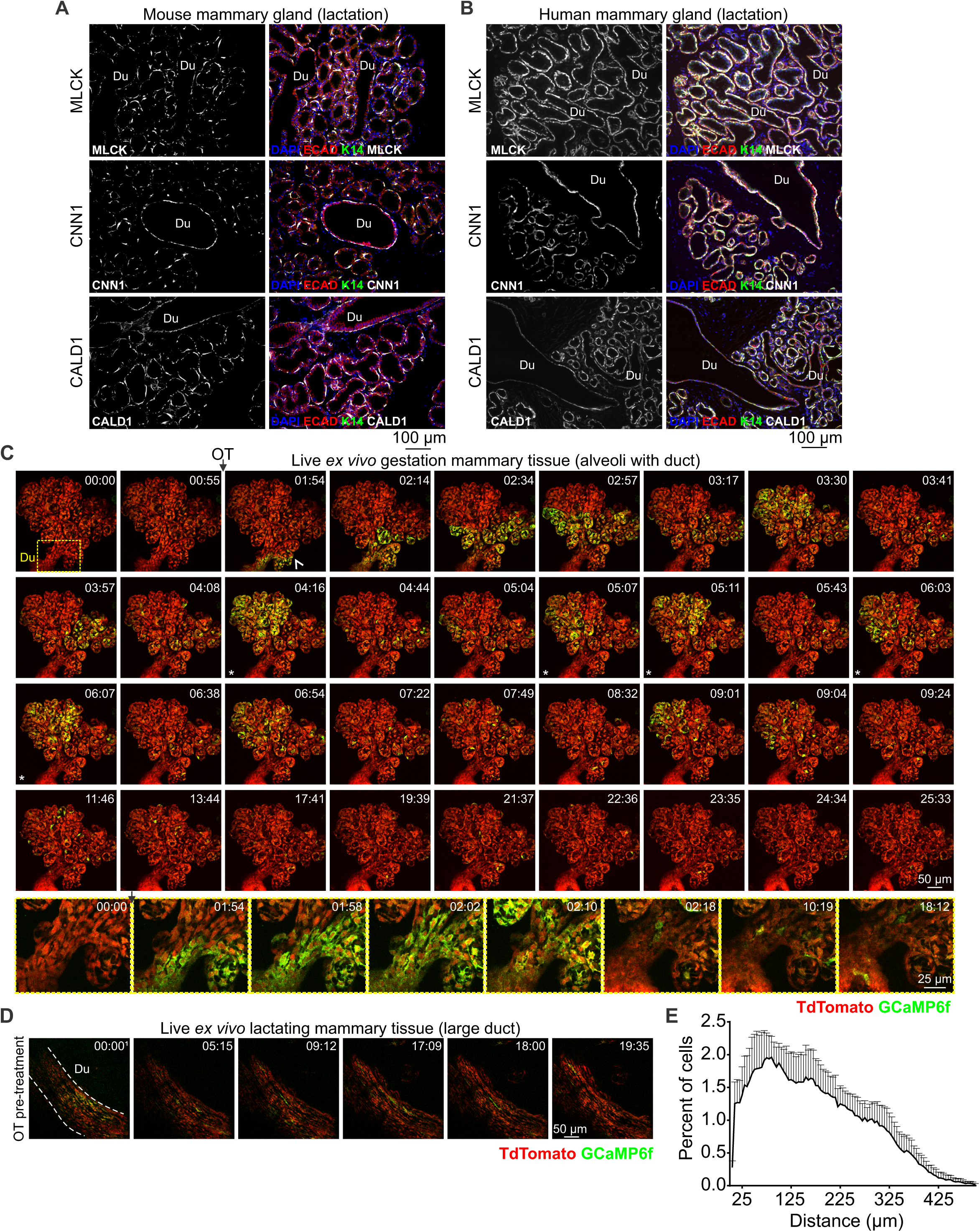
Functional differentiation and Ca^2+^-contraction coupling in ducts and alveoli. (**A-B**) Immunostaining of mouse and human lactating tissue. MLCK, CNN1 and CALD1 are expressed in both ducts (Du) and alveoli. E-cadherin (red) shows the luminal cell lineage; K14 (green) shows the basal cell lineage. Nuclei are stained with DAPI (blue); n = 3 samples, mouse and human. (**C**) 3D time-lapse imaging of live mammary tissue from a pregnant (15.5-16.5 d.p.c.) *GCaMP6f-TdTom;K5CreERT2* mouse stimulated with OT (85 nM) at 01:15 (min:s). Images show maximum intensity z-projection of 45 μm of live tissue; box (frame 1) shows subtending duct (Du, magnified in bottom panel), extending deeper into the tissue. Arrowhead at 01:54 shows direction of OT diffusion; asterisks show coordinated firing; n = 3. See also **Movie S6**. (**D**) 3D time-lapse imaging of a large duct from a lactating *GCaMP6f-TdTom;K5CreERT2* mouse stimulated with OT (85 nM) immediately prior to imaging. Images show maximum intensity z-projection of 45 μm of live tissue; n = 3. See also **Movie S7**. (**E**) Percent of cells with a high correlation coefficient (> 0.5) in Ca^2+^ firing and the Euclidean distance from correlated events. Graph shows average ± SEM (n = 4 mice, gestation).

Next, we used our model to examine possible Ca^2+^-contraction coupling in ductal cells of pregnant *GCaMP6f-TdTom;K5CreERT2* mice. At this developmental stage, contractile proteins are upregulated (Fig. S6C), Ca^2+^-contraction coupling is observed in alveolar structures (Movies S4-5) and the visualization of ducts is not encumbered by light scattering and/or absorptive properties of interposing structures. Ca^2+^-contraction coupling was clearly observed in ductal basal cells (Fig. 3C and **Movie S6**). On several occasions, large ducts that were positioned deep within the mammary tissue of lactating animals, were able to be observed (Fig. 3D and **Movie S7**), confirming these findings in the fully mature state. In mammary ducts, basal cells adopt a spindle-like morphology and are collectively oriented along the length of the duct (Fig. 1A). Our data reveal that contraction of ductal basal cells generates longitudinal motion, facilitating the continued flow of milk. We also demonstrate that differences in the type of motion generated by ductal and alveolar contractions arise from organizational heterogeneity—rather than divergent functional differentiation or signal transduction.

### Mammary epithelial cells *in situ* exhibit both stochastic and coordinated behaviors

Our model enables us to visualize molecular events in single cells, to observe how these events control an individual cell’s behavior and to understand how individual behaviors produce tissue-level outcomes. In mammary tissue, basal epithelial cells primarily exhibit stochastic activity (Figs 1-2 and Figs S4-5). Individual oscillatory behavior, however, was observed to be temporarily entrained across large lobuloalveolar structures (Fig. 3C, Figs S3-4 asterisks and Movies S2, S4 and S6), suggesting that this organ can generate both synchronized and unsynchronized motion for optimal milk ejection. To determine the degree of lobuloalveolar cooperativity in firing, we employed two agnostic approaches to analyze the functional connectivity in Ca^2+^ signaling events. First, we analyzed correlations in the firing pattern of individual basal cells in the post diffusion phase and graphed the Euclidean distances between highly correlated (> 0.5) cells. Highly correlated responses exhibited a short Euclidean distance (Fig. 3E). Next, we analyzed network topologies by connecting highly correlated cells within a single field-of-view. This method confirmed high clustering associated with short internodal distances in some lobular structures (small worldness) (Fig. S7) (24, 25). These analyses suggest some cooperativity in firing and, by extension, contraction.

### Distinct signaling pathways underpin the passage of milk, tears and sperm

To assess potential conservation in the signaling pathways that operate in basal cells of other OT-sensitive, fluid transporting epithelia, we assessed OT-mediated responses in the lacrimal glands and epididymides of *GCaMP6f-TdTom;K5CreERT2* mice. In the lacrimal gland, basal cells have a similar morphology, arrangement and function to mammary basal cells (26). They have previously been shown to undergo OT-dependent contractions (27), and diminished OT-OXTR signaling in these cells has been linked to dry eye disease (27). Like the mammary gland, dual expression of basal and smooth muscle markers was confirmed in lacrimal acini (Fig. 4A), however, no OT-mediated [Ca^2+^]_i_ or contractile responses were detected in these cells in this study (Fig. 4B-C and **Movie S8**).

**Figure 4.**
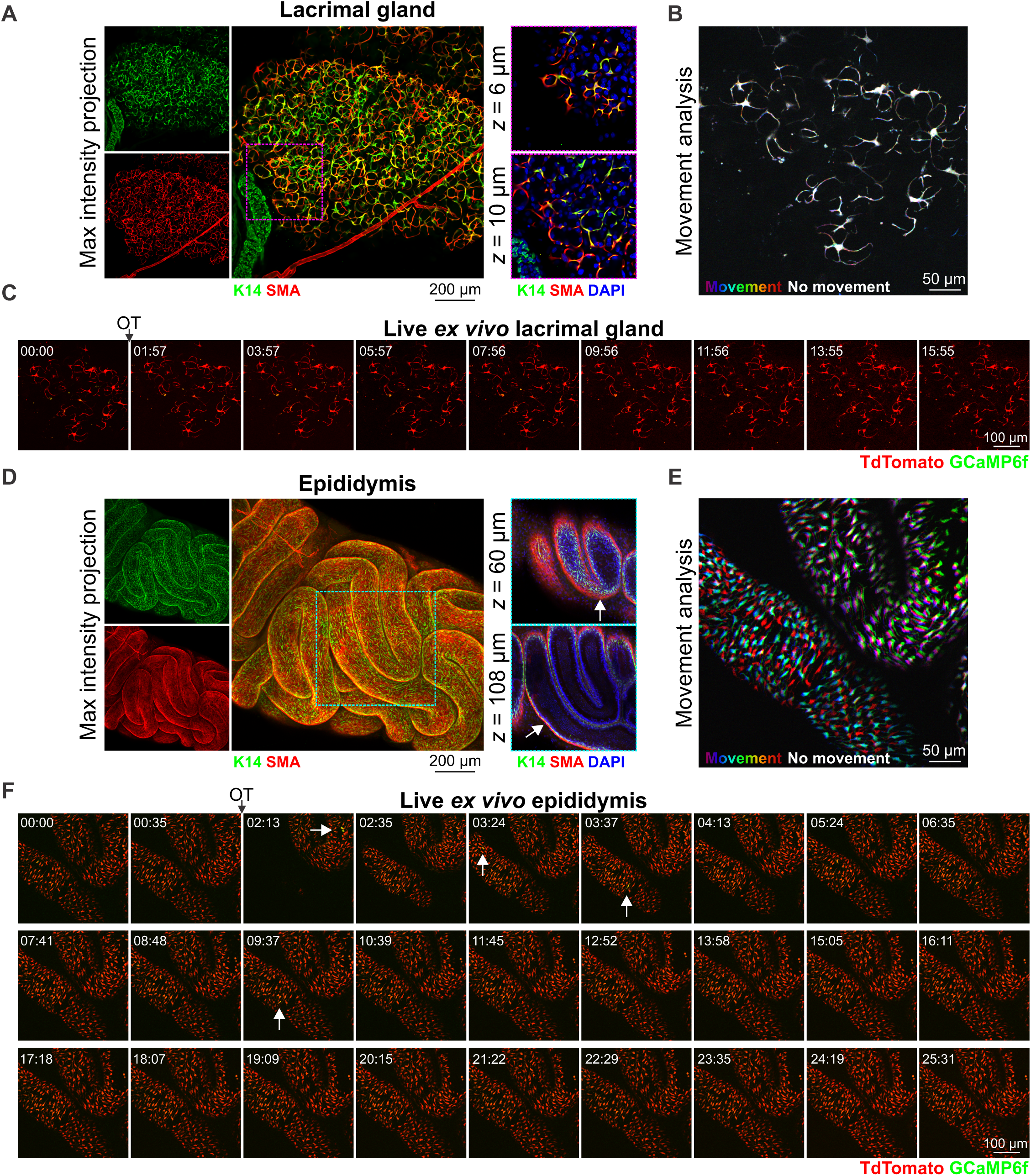
OT responses in basal epithelial cells of other fluid moving organs. (**A**) Maximum intensity z-projection (0-36 μm) and optical slices of lacrimal tissue. Lacrimal acinar basal cells express K14 and SMA. (**B**) Analysis of tissue movement created by the overlay of 3 images (each 43 s apart). Each image has been assigned a primary color (R-G-B). Regions that do not move during the 90 s window have R-G-B pixels superimposed and are white. Regions where significant movement has occurred appear R, G, B or a combination of 2 colors. See also **Movie S8**. (**C**) 3D time-lapse imaging of lacrimal tissue from *GCaMP6f-TdTom;K5CreERT2* mice. Tissue was stimulated with OT (85 nM, 00:45). Image series show maximum intensity z-projection (21 μm). (**D**) Maximum intensity z-projection (0-246 μm) and optical slices of cleared mouse epididymis (caput). Basal K14 positive cells are surrounded by SMA positive cells (arrow). (**E**) Tissue movement analysis of 3 images (45 s apart) as per (B). (F) 3D time-lapse imaging of epididymal tissue from *GCaMP6f-TdTom;K5CreERT2* mice. Tissue was stimulated with OT (850 nM, 01:38); arrows show single cell calcium responses. Series is maximum intensity z-projection 104.5 μm through the structure. See also **Movie S9**. N = 3 mice.

In males, a large burst of OT is released into the bloodstream at ejaculation (8, 28). This produces contractions of the male reproductive tract and, by assisting with the passage of fluid along this tract, these contractions are thought to reduce post-ejaculatory refractoriness and improve reproductive readiness (28, 29). Epididymal basal cells express basal cell markers, however, unlike the lacrimal and mammary glands, they do not co-express smooth muscle markers (Fig. 4D). Instead, movement of fluid through this organ appears to rely on a layer of smooth muscle surrounding the inner tubular epithelium (Fig. 4D). To assess the transport of sperm through this organ, its OT-responsiveness and its relationship to basal cell [Ca^2+^]_i_ elevations, we stimulated acute epididymal tissue pieces with a large bolus dose of OT. OT stimulation triggered marked peristaltic-like movements of the epididymal tubes (Fig. 4E) and a supra-basal pattern of phosphorylation of MLC (Fig. S8). Low frequency Ca^2+^ firing in basal cells was observed before and after OT-stimulation (Fig. 4F and **Movie S9**). Basal cell Ca^2+^-contraction signaling can therefore be selectively uncoupled in different fluid moving epithelia.

### Pharmacological inhibitors of regulatory proteins of myosin light chain phosphorylation are unable to block mammary contractions

Mammary basal cells express smooth muscle actin (Fig. S1A) and strongly upregulate elements of the vascular smooth muscle contraction pathway during gestation and early lactation (Fig. S6). Our group and others have therefore hypothesized that basal cell contraction is principally controlled by Ca^2+^/calmodulin-dependent phosphorylation of the myosin light chain (MLC) by MLCK and subsequent de-phosphorylation by myosin light chain phosphatase (MLCP) (9, 10, 30). This hypothesis is supported in the current study by a pattern of pMLC immunostaining in OT-treated tissue that is consistent with the organization of its Ca^2+^ firing activity (Fig. 1C-D and Fig. S3). To explore this further, we treated uterine, bladder, epididymal and mammary acute tissue pieces with pharmacological inhibitors of both MLCK and the MLCP inhibitor rho-associated protein kinase (ROCK) (Fig. S9A). Inhibition of MLCK and ROCK modestly reduced the intensity of contraction in all organs examined, although considerable variability was observed both within and between tissues (Fig. 5 and **Movie S10**). This is in contrast to a previous study, which scored contraction based on basal cell morphology in tissue treated prior to fixation with this ROCKi (31). When tissue was incubated with a cocktail of pharmacological inhibitors against MLCK, ROCK, protein kinase C (PKC) (32) and Ca^2+^/calmodulin-dependent protein kinase II (CaMKII) (33), contraction was robustly inhibited in uterine, epididymal and bladder preparations, but persisted in the mammary gland (Fig. 5A-B and **Movie S10**).

**Figure 5.**
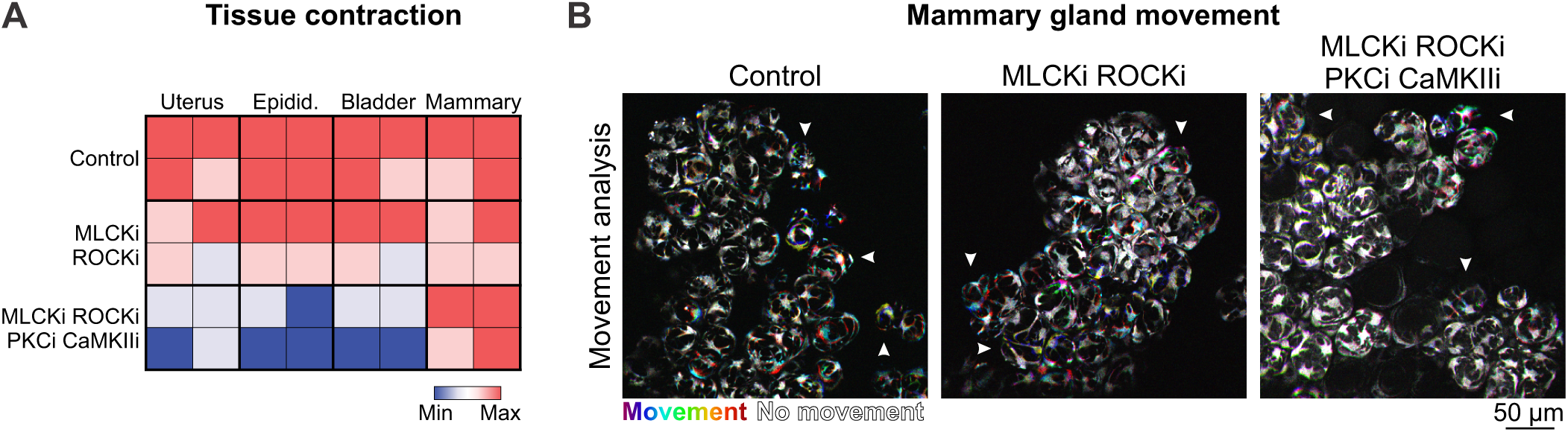
Pharmacological inhibition of the contractile pathway. (**A**) Matrix of contractile activity in acute tissue pieces isolated from uterus, epididymis, bladder and mammary gland and treated with either physiological buffer (control), a combination of inhibitors of MLCK (ML-9) and ROCK (Y27632) or a combination of inhibitors of MLCK (ML-9), ROCK (Y27632), PKC (calphostin-C) and CaMKII (KN93). Contractions were induced with oxytocin (85 nM, uterus and mammary gland; 850 nM epididymis) or carbachol (10 μM, bladder). See also **Movie S10**. (**B**) Analysis of tissue movement in acute mammary tissue pieces created by the overlay of 3 images (each 30 s apart). Each image has been assigned a primary color (R-G-B). Regions that do not move during the 90 s window have R-G-B pixels superimposed and are white. Regions where significant movement has occurred appear R, G, B or a combination of 2 colors. N = 4 mice.

It is conceivable that some pharmacological inhibitors are unable to effectively and consistently bind to their intracellular targets when applied to intact, lipid-rich mammary tissue. We therefore interrogated Ca^2+^-contraction coupling in dissociated primary mammary basal cells in a 2D assay. Cells from pregnant *GCaMP6f-TdTom;K5CreERT2* mice were isolated, plated in co-culture on a nanopatterned surface (Fig. S9B) and imaged within 12 h of dissection. These conditions were optimal for: 1) maintaining cell health and stage-specific differentiation; and 2) achieving anisotropy in the arrangement of contractile elements for the experimental measurement of force generation along a single axis (34). Under these conditions, OT stimulation produced [Ca^2+^]_i_ responses, which were coupled to contraction at the first (InsP3) phase (Fig. S9C shading and **Movie S11**). Later phase Ca^2+^-contraction coupling, however, was not able to be assessed in this model, due to the intensity of the first contraction (even at pM concentrations of OT) and the relatively low strength of the newly formed surface adhesions (12). Nevertheless, as Ca^2+^-contraction coupling is observed at this phase (Fig. 2D), we proceeded to use this system to examine this initial event in primary cells.

Intracellular Ca^2+^ chelation with BAPTA completely blocked [Ca^2+^]_i_ responses to OT (Fig. S9D and **Movie S12**). Cell contractions were also attenuated demonstrating, unequivocally, their Ca^2+^-dependence. To gauge the distance between the Ca^2+^ source (in this case InsP3 receptors) and sensor, we compared OT-mediated basal cell contractions in cells loaded with two different [Ca^2+^]_i_ chelators (BAPTA-AM and EGTA-AM), with different Ca^2+^ binding rates but comparable binding affinities (35, 36). Both intracellular BAPTA and EGTA were able to capture Ca^2+^ between the channel and the sensor (Fig. S9D), suggestive of “loose” Ca^2+^-contraction coupling in these cells that is not strictly dependent on nanodomain signaling (where EGTA is ineffective) (36). Similar to whole tissue preparations, however, treatment of cells with MLCK and ROCK inhibitors failed to block OT-mediated basal cell contraction (Fig. S9D). These data are not dissimilar to previous studies, where *in vitro* contraction was inhibited by only 30% in basal cells isolated from mice deficient for the gene encoding smooth muscle actin (30) and support a level of functional redundancy in the mammary contraction pathway.

### Coupled oscillator-based synchronization in the mammary gland

Ca^2+^-activation mechanisms in smooth muscle cells are incredibly diverse and are uniquely adapted to match the developmental stage-specific function of the biological structure on which they exert their force. Additional complexity arises when the mechanisms responsible for generating and propagating [Ca^2+^]_i_ signals in “smooth muscle-like” (myo)epithelial lineages are considered. Here, we demonstrate in mammary basal cells that OXTR engagement produces initial release of Ca^2+^ from intracellular stores, sufficient to generate cell and tissue contraction (Fig. 2D-E). Initial [Ca^2+^]_i_ responses have been shown to be sensitive to PLC inhibition in *in vitro* assays (12) and similar [Ca^2+^]_i_ responses are observed with InsP3 infusion (12), consistent with coupling via Gq-proteins to PLCβ (37). In some smooth muscle cells, [Ca^2+^]_i_ signals are propagated along the length of the cell via the regenerative release of stored Ca^2+^ by ryanodine receptors (RYRs) (38, 39). As cytosolic Ca^2+^ waves were also observed in mammary basal cells (Fig. 2A), we investigated novel roles for RYRs in this tissue. *Ryr1* (but not -*2* or -*3*) was expressed in lysates that were prepared from homogenized mammary tissue during lactation (Fig. S10A) and was enriched in functionally mature basal cells (Fig. S10B). To determine the role of RYR1 channels in these cells, we treated mammary tissue from *GCaMP6f-TdTom;K5CreERT2* mice with the ryanodine receptor inhibitor dantrolene (40). Dantrolene did not inhibit the initial release of Ca^2+^ from intracellular stores (Fig. 6A and **Movie S13**). However, to our surprise, [Ca^2+^]_i_ oscillations became entrained in some regions and tissue exhibited rhythmic and sustained pulses of activity that resembled smooth muscle phase waves, with a periodicity of 104.2 ± 16.38 s and a velocity of 10.62 ± 2.64 μm.s^-1^ (Fig. 6A-B and **Movie S13**). A similar effect was observed with inhibiting concentrations of the plant alkaloid ryanodine (41) (**Movie S14**). These data, together with our observation that [Ca^2+^]_i_ oscillations could be temporarily entrained under physiological conditions (Fig. 3C, Fig. S4 and Movies S2 and S6), support a model whereby mammary basal cells can alternate between unsynchronized movements and coupled oscillator-based lobuloalveolar synchronization, modulated in-part by the mechanism of ER Ca^2+^ release.

**Figure 6.**
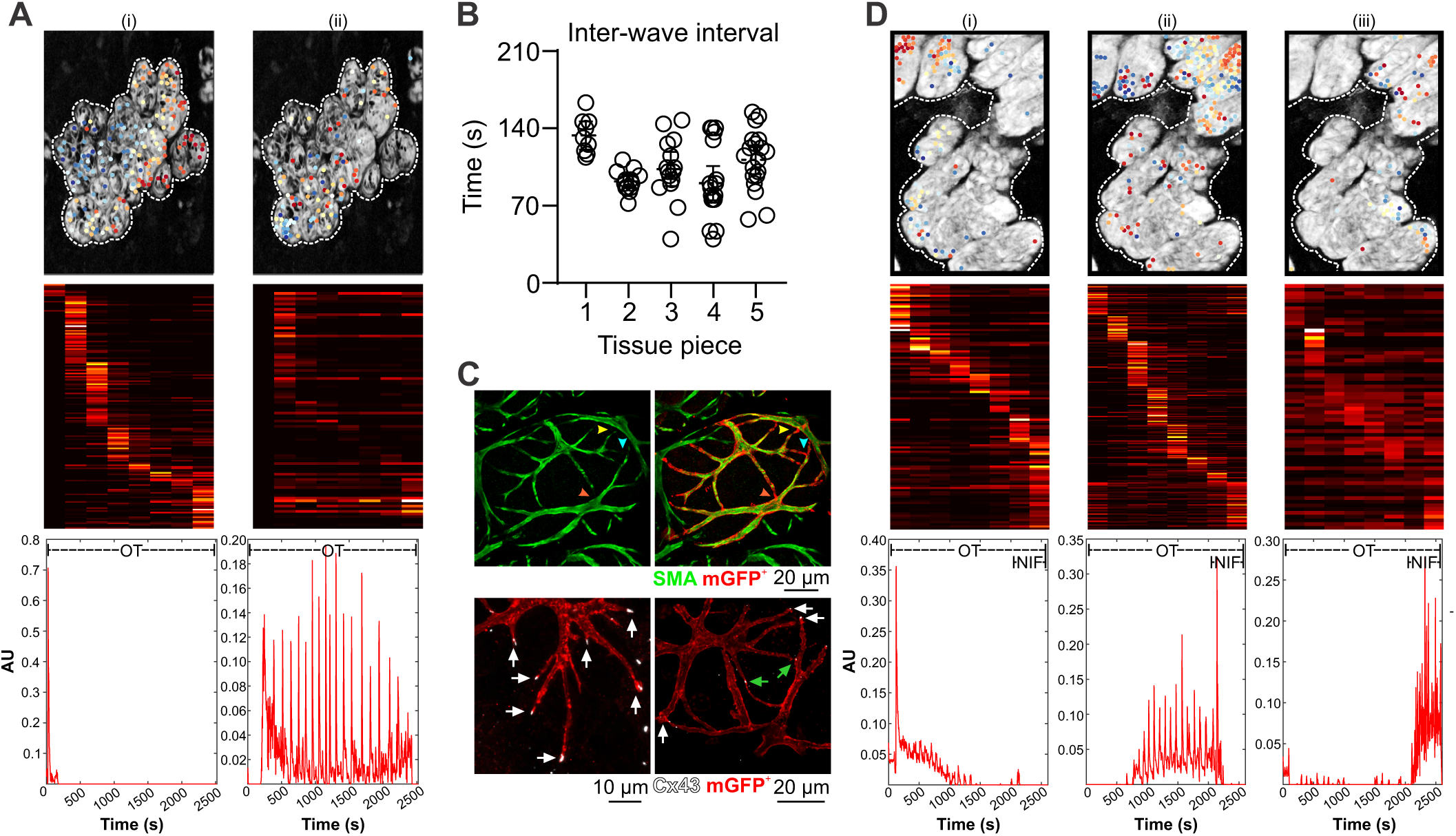
Dantrolene-induced tissue synchronization. (**A**) Sequential Non-Negative Matrix Factorization (seqNMF) was used to identify sequences of events and cluster cells functionally. (i) corresponds to the initial InsP3 response, (ii) corresponds to the dantrolene dependent synchronized activity. Dots (top panel) are color-coded according to their temporal position in the sequence of events (middle panel) and overlaid on a maximum intensity z-projection of the green channel. The sequence of [Ca^2+^]_i_ activity within each cluster is represented in the bottom panel; n = 3 mice. (**B**) Interval between each wave of synchronized activity in ex vivo dantrolene-treated mammary tissue (mean +/- 95% CI); n = 5 tissue pieces from at least 3 mice. (**C**) Optically-cleared mammary tissue from lactating mice showing SMA immunostaining (green, top panel) and two cells expressing a membrane targeted fluorescent protein (red, top panel). Colored arrowheads point to sites of cell-cell contact that are revealed by the membrane marker. Immunostaining for Cx43 (white, bottom panel) in cells expressing membrane targeted fluorescent protein (red, bottom panel). White arrows show Cx43 staining at basal cell tips; green arrows show Cx43 staining at sites where two labelled basal cells are connected. Membrane marker has been induced at near clonal density (i.e. does not stain all basal cells); n = 3 mice. (**D**) seqNMF as in A, (i) corresponds to the initial InsP3 response, (ii) corresponds to the dantrolene dependent synchronized activity, (iii) corresponds to the addition of nifedipine to the mammary tissue. After addition of nifedipine, the synchronized activity disappears and switches to a stochastic activity distributed through the tissue. See also **Movie S15**. AU, arbitrary unit; n = 3 mice.

A key factor of coupled oscillator-based synchronization is intercellular communication via gap junctions (38, 42). Mammary basal cells express Cx43 (43, 44) and mice with severely compromised Cx43 function have impaired milk ejection (45). However, it is often difficult to appreciate how stellate basal cells are physically coupled to their neighbors when visualized using thin tissue sections (Fig. S11). Similarly, due to their size and exclusion from near plasma membrane domains, the true extent of basal cell connectivity has not yet been captured using 3-dimensional imaging of conventional basal cell markers (Fig. S1A). To overcome this, we developed mice that express a membrane localized fluorescent protein in basal cells and assessed Cx43 localization in optically cleared tissue. Using this approach, basal cell boundaries were readily identified, enabling us to visualize how thin processes of adjacent cells are physically connected (Fig. 6C, top panel). Cx43 was enriched at sites of homotypic cell contact (Fig. 6C, bottom panel). These data confirm that the cytoplasms of adjacent basal cells are linked, enabling individual cells to coordinate the activity of the larger system.

In other tissue types that exhibit rhythmic contractions, e.g., vascular, lymphatic and airway smooth muscle, periodic release of Ca^2+^ from the ER produces membrane depolarization and activation of L-type Ca^2+^ channels (38). Current flow through gap junctions enables depolarization to spread rapidly into neighboring cells, synchronizing large numbers of cells potentially over millimeter distances (38, 42). To determine whether L-type calcium channels are involved in synchronization events in the mammary gland, we treated rhythmically contracting tissue with the L-type Ca^2+^ channel blocker nifedipine. Nifedipine rapidly and consistently resulted in the reversion to stochastic activity (Fig. 6D and **Movie S15**). Collectively, these data reveal that mammary basal cells are physically and electrically coupled, enabling Ca^2+^ to control both the behavior of individual cells as well as the system as a whole.

## Discussion

Real-time, *in situ* activity monitoring provides important insights into how individual cells behave in multi-dimensional and multi-cellular environments (1–4). This approach was used to describe and quantify the mechanism by which milk is transported through the hollow mammary epithelium, making it available on-demand and with minimal delay to the nursing neonate (8, 46). Our data support a number of novel conclusions that could not have been obtained using conventional methods.

Firstly, we revealed that transient [Ca^2+^]_i_ elevations precede and are required for basal cell contractions in the functionally-mature gland. We extended this finding to demonstrate how Ca^2+^-contraction coupling in a single basal cell can physically warp the layer of alveolar luminal cells that it encircles. Structure, function and expression were examined in the adjoining ductal epithelium, previously relegated to a role akin to a biological drinking straw. Instead, our analyses revealed active participation of the ductal epithelium in the process of milk ejection. Differences in the type of motion generated by basal cell contractions in ducts and alveoli were ascribed to heterogeneity in cellular organization, rather than expression or function of contractile elements.

We explored components of the contractile network downstream of Ca^2+^ activation in mammary basal cells. A pattern of pMLC positivity was observed in mammary cell ensembles, which mirrored the Ca^2+^ activity of the tissue. Pharmacological inhibition of the Ca^2+^-dependent MLCK and the Ca^2+^-sensitizer ROCK, however, failed to block mammary contractions in our study. Whilst MLCK is widely considered to be the primary Ca^2+^-dependent regulator of MLC phosphorylation in smooth muscle, this model is based on reductionist principles, does not fit all smooth muscle cell types and fails to acknowledge the growing complexity in regulatory kinases known or hypothesized to govern smooth muscle contraction *in vivo* (47–49). Indeed, embryonic blood vessels from MLCK knockout mice remain responsive to cytosolic Ca^2+^ elevations (50). Our data reveal that, similar to aortic smooth muscle cells, “smooth muscle-like” epithelial cells in the mammary gland also display considerable complexity and diversity in their biomechanical behavior. Complexity in the pathways downstream of Ca^2+^ activation may extend beyond Ca^2+^-contraction coupling to Ca^2+^-transcription coupling (51), an aspect of signaling that has not been considered here but which may be relevant for the interpretation of genetic knockout models (9).

In addition to the diversity in signal transduction downstream of Ca^2+^ activation in mammary basal cells, our study and others (9, 12, 52) have demonstrated that a number of Ca^2+^ channels—with distinct activation mechanisms and cellular localizations—participate in its encoding. These include channels that regulate Ca^2+^ release from intracellular stores, influx from the extracellular environment and movement between the cytosol of adjacent cells. In this sense, [Ca^2+^]_i_ acts as a central node in a type of bow-tie motif in basal cells (53), whereby multiplicity in its encoding and decoding enable this evolutionarily essential organ to engage local and global motions to ensure adequate nutrition for the dependent offspring, while on-the-other-hand remaining vulnerable at this crucial point of convergence.

The dynamic nature of the oscillatory Ca^2+^ signal enables basal cells to rapidly cycle between contracted and relaxed states. We posit that the spatiotemporal properties of this signal are important insomuch as its oscillation intensity and interval match the activation threshold and decay rate of the downstream effector to permit efficient switching between cycles of contraction and relaxation. Coupling of the Ca^2+^ sensor within nanometer distance to the channel pore, however, appears unlikely based on the following new observations: 1) both ER Ca^2+^ release and plasmalemmal Ca^2+^ influx were sufficient for *in situ* basal cells to develop and bear tension; and 2) BAPTA-AM and EGTA-AM were equally effective in inhibiting *in vitro* contractions, despite EGTA’s slower binding kinetics. Although not essential for Ca^2+^-contraction coupling, highly spatially regulated [Ca^2+^]_i_ signals may be an important factor for Ca^2+^-transcription coupling for the long-term maintenance of the contractile phenotype (51) or Ca^2+^ wave generation at the tissue-level.

Finally, our data, together with published work, suggests that mammary basal cells are able to shift between store- (9) and voltage-dependent modes of operation, a phenomenon that appears to be moderated, at least in-part, by the mechanism of ER Ca^2+^ release. It is currently unclear how basal cells coordinate the activity of these two, often reciprocally regulated (54, 55), influx pathways under physiological conditions. However, our observation that pharmacological inhibition of RYR1 promoted dihydropyridine-sensitive signal synchronization, corresponds with accounts of RYR activity in bona fide smooth muscle cells (51). Here, RYR-mediated Ca^2+^ sparks can activate nearby BK_Ca_ channels, producing spontaneous transient outward currents (STOCs), membrane hyperpolarization and reduced Cav1.2 activity (38, 51). Optical monitoring of voltage in 3-dimensions using genetically-encoded voltage indicators (GEVIs) (56) and examination of population dynamics in *Cacna1c*-, *Ryr1*- and *Kcnma1*-conditional knockout mice remain aims for the future. It is also unclear at this time whether spatial synchronicity can be initiated by any oscillating basal cell (alveolar or ductal) within the mammary epithelium or whether basal cells lock into the frequency of a putative population of epithelial (57) or interstitial (58) mammary “pacemaker” cells. This question may be addressed by future studies using light-sheet fluorescence microscopy and quantitative image analysis to create a spatial footprint of the frequency dynamics of individual oscillators and phase advanced cells.

In summary, by imaging activity in the mammary gland across scales, we were able to visualize and describe in unprecedented detail how the repetitive and collective effort of thousands of mammary basal cells facilitate the transport of a thick biological emulsion through a narrow passage in a manner that is both consistent and persistent. Moreover, the system presented here represents a novel, physiologically relevant model for studying the collective nature of mammalian biological processes.

## Materials and Methods

### Mice

Animal experimentation was carried out in accordance with the *Australian Code for the Care and Use of Animals for Scientific Purposes and the Queensland Animal Care and Protection Act* (*2001*), with local animal ethics committee approval. Animals were kept in a Specific Pathogen Free facility under a 12:12 h light-dark cycle in individually ventilated cages. Food and water were available ad *libitum* and standard enrichment provided. All strains were purchased and maintained on a C57BL6 background. *K5CreERT2* mice (B6N.129S6(Cg)-Krt5^tm1.1(cre/ERT2)Blh^/J, stock no. 029155), *Lck-GCaMP6f-fl* mice (C57BL/6N-Gt(ROSA)26Sor^tm1(CAG-GCaMP6f)Khakh^/J, stock no. 029626) and Acta2-Cre mice (B6.FVB-Tg(Acta2-cre)1Rkl/J, stock no. 029925) were purchased from The Jackson Laboratory (Bar Harbor, ME). TdTomato-fl mice (B6.Cg-Gt(ROSA)26Sor^tm9(CAG-tdTomato)Hze^/J, stock no. 007909) were a kind gift from Prof. Ian Frazer (University of Queensland). *GCaMP6f-fl* mice (B6J.Cg-*Gt(ROSA)26Sor^tm95.1(CAG-GCaMP6f)Hze^*/MwarJ) were a kind gift from Dr James W. Putney Jr (National Institute of Environmental Health Sciences). C57BL6/J mice were obtained from the Animal Resources Centre (Western Australia). All Cre-expressing lines were maintained as heterozygotes.

Genotyping was performed on mouse toe or ear DNA using the following primers: to distinguish K5-CreERT2 5’-GCA AGA CCC TGG TCC TCA C-3’, 5’-GGA GGA AGT CAG AAC CAG GAC-3’, 5’-ACC GGC CTT ATT CCA AGC-3’ (wildtype 322 bp, mutant 190 bp); to distinguish *TdTomato-fl*, *GCaMP6f-fl* and *Lck*-*GCaMP6f-fl* 5’-CTC TGC TGC CTC CTG GCT TCT-3’, 5’-CGA GGC GGA TCA CAA GCA ATA-3’ and 5’-TCA ATG GGC GGG GGT CGT T-3’ (wildtype 330 bp, mutant 250 bp); and to distinguish Acta2-Cre 5’ACA TGT CCA TCA GGT TCT TGC-3’, 5’-AGT GGC CTC TTC CAG AAA TG-3’, 5’-TGC GAC TGT GTC TGA TTT CC-3’ and 5’-GGT GTT AGT TGA GAA CTG TGG AG-3’ (wildtype 521 bp, mutant 300 bp).

To induce expression of GCaMP6f and TdTomato in K5-positive cells, female mice were administered tamoxifen (1.5 mg) (Sigma Aldich, T5648) diluted in sunflower oil (Sigma Aldrich, S5007) and ethanol (10%) via intraperitoneal injection at 4-weeks of age. A further 2-3 tamoxifen injections were administered every second day on alternating sides at 8-weeks of age, providing a total dose of 4.5-6 mg per mouse. A 4-6-week washout period was observed before mating. Male mice were injected with 4 × 1.5 mg tamoxifen injections every second day at 8-weeks of age (total dose 6 mg per mouse). Female experimental mice were mated in pairs or trios with wildtype sires. To obtain mammary tissue during gestation, sires were removed after observation of a copulation plug and mammary tissue harvested 15.5-16.5 d.p.c (days post coitus). To obtain lactating tissue, sires were removed prior to littering and female mice were allowed to nurse for 8-14 days (peak lactation).

### Human subjects

Healthy tissue biopsies from consented lactating women (25-33 years old) were obtained from the Susan G. Komen Tissue Bank at the IU Simon Cancer Center (59). Tissue donors were recruited under a protocol approved by the Indiana University Institutional Review Board (IRB protocol number 1011003097) and according to The Code of Ethics of the World Medical Association (Declaration of Helsinki), with site-specific approval from the Mater Misericordiae Ltd Human Research Ethics Committee. Breast biopsies were fixed in formalin and paraffin embedded as per standard protocols. Lactating samples were collected from women who were actively breastfeeding at the time of tissue donation (at least once per day). Donors included in this study had been breastfeeding for 6 to 23 months prior to tissue donation.

### Immunohistochemistry

IHC was performed based on a previously published protocol (60). Briefly, formalin-fixed paraffin embedded mouse and human slides were deparaffinized in xylene and rehydrated in a reducing ethanol series. Tissue was permeabilized in phosphate buffered saline (PBS) containing triton X-100 (0.5%). Heat-induced epitope retrieval was performed in sodium citrate (0.01 M, pH 6) for 11 min at 110°C using a NxGen Decloaking Chamber. Slides were blocked in PBS containing normal goat serum (10%) and triton X-100 (0.05%) for 1 h. Primary antibodies were incubated overnight at 4°C in a humidified chamber. The following primary antibodies were used in this study: rabbit anti-SMA (Abcam, ab5694, 1:600), rabbit anti-MLCK (Sigma Aldrich, HPA031677, 1:700-1:1000), rabbit anti-CNN1 (Abcam, ab46794, 1:700-1:1000), rabbit anti-CALD1 (Sigma Aldrich, HPA008066, 1:200), mouse anti-E-cadherin (BD Biosciences, 610182, 1:200-1:400), chicken anti-K14 (BioLegend, 906004, 1:200-1:400) and rabbit anti-Cx43 (Abcam, ab11370, 1:1000). Secondary antibodies were incubated for 1 h at room temperature. The following secondary antibodies were used in this study (1:500): goat anti-rabbit Alexa Fluor 647 (ThermoFisher Scientific, A21236), goat anti-mouse Alexa Fluor 555 (ThermoFisher Scientific, A32727) and goat anti-chicken Alexa Fluor 488 (ThermoFisher Scientific, A11039). Nuclei were stained with DAPI dilactate (625 ng/mL) for 10 min at room temperature and tissue was imaged on an Olympus BX63F upright epifluorescence microscope.

### Tissue clearing

Mammary tissue was dissected, spread on foam biopsy pads and optimally fixed in neutral buffered formalin (NBF, 10%) for 3-9 h (according to size) at room temperature. Lacrimal and epididymal tissue was dissected and fixed in NBF for 4-7 h. Pre-stimulation with OT [85 nM (lacrimal and mammary), 850 nM (epididymis), 2-10 min] was performed on live tissue prior to fixation, as indicated. Tissue clearing was performed on fixed tissue using a modified CUBIC protocol (16, 61), except for studies examining Cx43 and the membrane targeted fluorescent protein, which used SeeDB (62). For CUBIC clearing, tissue was immersed in CUBIC Reagent 1A [urea (10% w/w), Quadrol® (5% w/w), triton X-100 (10% w/w), NaCl (25 mM) in distilled water] for 2-3 days at 37°C, washed in PBS and blocked overnight at 4°C in PBS containing normal goat serum (10%) and triton X-100 (0.5%). Tissue was incubated in primary antibodies diluted blocking buffer at 4°C for 4 days. The following primary antibodies and dilutions were used for wholemount immunostaining in this study: rabbit anti-SMA (1:300), chicken anti-K14 (1:200), rabbit anti-pMLC (Abcam, ab2480, 1:200) and chicken anti-GFP (Abcam, ab13970, 1:1000-1:2000). Secondary antibodies (Life Technologies) are listed above and used 1:500, incubated for 2 days at 4°C for wholemount immunostaining. Nuclei were stained with DAPI dilactate (5 μg/mL) for 2-3 h at room temperature. Washing was performed in PBS after primary and secondary antibody incubations (3 × 1 h). Tissue was placed in modified CUBIC Reagent 2 [sucrose (44% w/w), urea (22% w/w), triethanolamine (9% w/w), triton X-100 (0.1% v/w) in distilled water] at 37°C and imaged within 1-7 days (61, 63). Tissue was imaged in CUBIC Reagent 2 using an Olympus FV3000 laser scanning confocal microscope with UPLSAPO 10×/0.4, UPLSAPO 20×/0.75 and UPLFLN 40×/0.75 objective lenses. For SeeDB clearing, tissue was blocked overnight at 4°C in PBS containing BSA (10%) and triton X-100 (1%) and incubated in primary antibodies diluted blocking buffer at 4°C for 4 days [chicken anti-GFP, rabbit anti-SMA or rabbit anti-Cx43 (1:200)]. Tissue was washed and incubated in secondary antibodies and DAPI as described above, before moving through increasing fructose solutions (16). Imaging depth (z) is recorded from the start of the image sequence, typically 50-150 μm through the cleared structure. Visualization and image processing was performed in Imaris (v9.2.1) or ImageJ (v1.52e, National Institutes of Health) (64, 65). Denoising of 3D image stacks was performed as previously described (66).

### Primary cell imaging

Abdominal and inguinal mammary glands were dissected from pregnant *GCaMP6f-TdTom;K5CreERT2* mice (15.5-16.5 d.p.c.), lymph nodes removed and mammary tissue finely chopped. Cells were isolated using adapted published protocols (67, 68). Briefly, diced tissue was incubated at 37°C with gentle rocking in DMEM/F-12 (with L-glutamine and HEPES, ThermoFisher Scientific 11330032) supplemented with collagenase (Roche, 11088793001, 1 mg/mL) and hyaluronidase (Sigma Aldrich, H3506, 100 U/mL). Cells were disrupted every 30 min using a transfer pipette. After 3 h the tube was removed from the incubator, cells were further disrupted by pipetting and pelleted (1500 rpm, 5 min, 4°C). The subsequent steps and washes were performed in Hank’s Buffered Salt Solution with fetal calf serum (2%). Red cell lysis was performed in ammonium chloride. Pelleted cells were treated with pre-warmed trypsin (0.25%) for 1 min. Re-suspended and re-pelleted cells were then treated with pre-warmed dispase (5 mg/mL) and DNase (1 mg/mL) for 1 min, re-suspended and strained through a 40 μm cell strainer. Cells were plated in DMEM/F12 with FCS (10%) on collagen coated (50 μg/mL) NanoSurface 96-well plates (NanoSurface Biomedical ANFS-0096) and allowed to attach for at least 2.5 h.

Cells were imaged in physiological salt solution [PSS; HEPES (10 mM), KCl (5.9 mM), MgCl2 (1.4 mM), NaH2PO4 (1.2 mM), NaHCO3 (5 mM) NaCl (140 mM), glucose (11.5 mM), CaCl2 (1.8 mM); pH 7.3-7.4] within 12 h from the time of sacrifice. Cells were loaded with BAPTA-AM (100 μM, ThermoFisher Scientific, B6769) and EGTA-AM (100 μM, ThermoFisher Scientific, E1219), as indicated, in complete media at 37°C for 20-40 min before imaging. Pre-treatments with ML-7 (20 μM, Abcam, Ab120848), Y27632 (10 μM, Abmole, M1817) or a combination of ML-7 and Y27632 were performed for 10-20 min in PSS. Cells were imaged on an Olympus FV3000 inverted confocal microscope using a UPLSAPO 30×/1.05 silicone objective, resonant scanner and z-drift compensator. The rise time (Tpeak) for GCaMP6f is approximately 80 ms; the half decay time (T1/2) is 400 ms. Images were acquired every 3 s (< firing duration) for 6 min. Cells were stimulated with OT (0.85 nM) at 30 s; the concentration of ML-7 and Y27632 were maintained for the duration of the assay.

### *Ex vivo* tissue imaging

Mammary glands and uteri were harvested from lactating wildtype, *GCaMP6f;K5CreERT2* or *GCaMP6f-TdTom;K5CreERT2* mice, diced into 3-4 mm^3^ pieces and loaded with CellTracker™ Red (1.5 μM) in complete media for at least 20 min at 37°C and 5% CO2, as indicated (9). Lacrimal glands (extraorbital), epididymides and bladders were isolated from wildtype or *GCaMP6f-TdTom;K5CreERT2* mice and loaded with CellTracker™ Red, as indicated. Cut (undigested) tissue pieces were anchored with netting and imaged in 35 mm imaging dishes, bathed in PSS. Tissue was stimulated with OT (85 nM, mammary, uterus and lacrimal; 850 nM epididymis) or carbachol (10 μM, bladder) in PSS, while imaging on an Olympus FV3000 LSM using a resonant scanner and UPLSAPO 30×/1.05 silicone objective. Images were acquired every 1.8-5 s for 15-45 min, depending on tissue type and experimental conditions.

For extracellular Ca^2+^ chelation and addback experiments, tissue was incubated in Ca^2+^-free PSS containing BAPTA (2 mM) for 40 min prior to imaging. Tissue was incubated in pharmacological inhibitors in complete media for 2-6 h prior to imaging. The following pharmacological inhibitors were used in this study: ML-9 (Tocris, 0431/10, 20 μM), Y27632 (Abmole, M1817, 20 μM), KN93 (Tocris, 12791, 20 μM), calphostin C (Tocris, 1626, 1 μM), dantrolene (Tocris, 0507, 20μM), ryanodine (Tocris, 1329, 100 μM), nifedipine (Sigma Aldrich, N7634, 20 μM). Concentrations of all chelators and inhibitors were maintained for the duration of the imaging study. Uterine, epididymal, bladder and mammary contractions were scored on a 4-point scale by a blinded investigator.

### Intravital imaging

Intravital imaging was performed on lactating mice (day 10-15) as previously described (19, 69). Nursing pups were removed and euthanized immediately prior to starting the experiment. The lactating dam was anesthetized using a combination of Zoletil® (tiletamine hydrochloride and zolazepam hydrochloride, 50 mg/kg) and xylazine (20 mg/kg) by intraperitoneal injection. Anesthesia was maintained using gaseous isofluorane (1.5%), supplied via a nose cone. Abdominal hair was removed using electronic pet clippers and Nair hair removal cream. An incision was made along the midline and hind limb to create a skin flap that extended from the abdominal midline to the spine and left the peritoneum intact. Skin was glued using Vetbond tissue adhesive to a standard microscope slide. Care was taken so that the tissue adhesive did not directly contact the mammary tissue. Vaseline and parafilm was used to protect the abdominal wall. A heat lamp was used to maintain body temperature. Blu Tack® was used to secure the slide-mounted skin flap onto the microscope stage above a #1.5 coverslip. Imaging was performed using an inverted Olympus FV3000 laser scanning microscope with a resonant scanner and UPLSAPO 30×/1.05 silicone objective. Before starting imaging, a top-up dose (30% induction dose) of injectable anesthetic was administered. Oxytocin (2.2 U) was administered via i/p injection approximately 1 min after the commencement of imaging. Vitals were monitored throughout the experiment, which was completed within 1 h from the induction of anesthesia.

### Registration and analysis of live 4D tissue imaging

We used Advanced Normalization Tools (ANTs, github.com/ANTsX/ANTs) to register the red fluorescence of the live Ca^2+^ imaging to the first time point of each movie (20, 21). We used the following call on each image of the image sequence : antsRegistration -d 2 --float 1 -o [OutImg, OutImg.nii] -n WelchWindowedSinc -w [0.005,0.995] -u 1 -r [FixImg,MovImg, 1] -t rigid[0.1] - m MI[FixImg,MovImg,1,32, Regular,0.25] -c [1000×500×200×50,1e-7,5] -f 8×4×2×1 -s 2×1×1×0vox -t Affine[0.1] -m MI[FixImg,MovImg,1,32, Regular,0.25] -c [1000×500×200×50,1e-7,5] -f 8×4×2×1 -s 2×1×1×0vox -t SyN[0.05,6,0.5] -m CC[FixImg,MovImg,1,2] -c [100×500×200×50,1e-7,5] -f 8×4×2×1 -s 2×2×1×0vox -v 1. We also computed the Jacobian determinant of the resulting warps, this provided an estimate of the “strength” of the warp required to align the two images, as such it is a proxy metric of contractions. Large movements and drifts would not be included in the Jacobian determinant as these are corrected by the rigid and affine step of the registration.

The resulting warps were then applied to the GCaMP6f channel and the resulting registered movies were analyzed using MATLAB. First, we manually selected regions of interests (ROIs) and extracted the fluorescence time series of each channel. The ΔF/F0 was then computed as described previously (70), and all the GCaMP6f peaks with at least a 10% increase in ΔF/F0 (change in starting fluorescence divided by starting fluorescence), and a correlation to an example GCaMP6f trace above 0.6, were identified. The time of those peaks was used to select a time period of five frames before and ten frames after the peak in both green and red channels, these time traces were then averaged across all identified peaks within and between each mice tissue. The delay between the peaks was also evaluated with the same criteria.

To uncover any potential coordination in the firing pattern, we used a correlation-based approach. We extracted the Ca^2+^ traces from live imaging of alveolar units using the CaImAn python package (71). A correlation distance matrix was then built between all the identified ROIs, as well as a Euclidean distance matrix between their centroids, for each of the mice analyzed. These were used to either look at the probability distribution of the Euclidean distance for highly correlated (> 0.5) ROIs, or to build a weighted directed graph between these ROIs.

The same approach was used to identify sequences of synchronized activity in the Dantrolene treated tissues, we used the sequential Non-negative Matrix Factorization (seqNMF) to cluster the Ca^2+^ traces in up to 3 clusters, with a regularization factor of 0.0001 and a time window of 8 (72). The number of clusters and regularization factor value were determined empirically to minimize the dissimilarity metric and optimize the reconstruction vs x-ortho cost, see Ref (72) for details. To estimate the time interval between each wave of synchronized activity, we measured the time between subsequent peaks of the repeated sequence identified by seqNMF. This approach was also used to measure the velocity, by dividing the Euclidean distance between active cells with the lag each cell presents in the sequence of activation.

### qPCR

For gene expression studies from whole tissue, snap frozen virgin and lactating mammary tissue (30-100 mg) was crushed under liquid nitrogen using a mortar and pestle, lysed in Buffer RLT Plus (Qiagen) and homogenized using QIAshredder homogenizer columns (Qiagen). For gene expression studies from sorted cells, primary cells were dissociated from pregnant C57BL/6J mice as described above and prepared for flow cytometry as previously described (67). Luminal and basal cells were identified by CD49f and EpCAM expression, after exclusion of non-epithelial cells and dead cells (67). RNA isolation and purification was performed using the RNeasy Plus Mini Kit (Qiagen) with gDNA eliminator columns and cDNA was prepared using the QuantiTect Reverse Transcription kit (Qiagen). Resulting cDNA was amplified using an ABI ViiA 7 real-time PCR System, using TaqMan Fast Universal PCR Master Mix and TaqMan Gene Expression Assays, with a limit of detection set as 36 cycles. The following Gene Expression Assays were used in this study: *Ryr1* (Mm01175211_m1), *Ryr2* (Mm00465877_m1), *Ryr3* (Mm01328395_m1), *Krt14* (Mm00516876_m1) and Esr1 (Mm00433149_m1). Relative quantification was calculated relative to 18S ribosomal RNA and analyzed using the comparative CT method (73).

### Gene expression analysis from published datasets

Preprocessed expression count data were obtained from Karsten Bach (74) and analyzed using scripts provided by Bach et al., modified to compare the C12 vs C14 clusters and C13 vs C14 clusters using edgeR 3.22.5 in R 3.5.1 (75). An R script to generate the differential expression results between these clusters is available at: (https://gist.github.com/adamewing/2819d7e5072aa35632bba7f51236446b). Violin and volcano plots were generated using the seaborn package for python (76); zero counts were removed. Scripts are available at: (https://gist.github.com/adamewing/931cc44d717959073fa5b09078e7e4b3) and (https://gist.github.com/adamewing/90f1f57985a2033f292debb2e2e5b25f) respectively. Gene Set Enrichment Analysis (GSEA) (77) was carried out using genes ranked by fold change.

### Primary cell contraction analysis

Images were quantified using a custom ImageJ macro for reproducibility and to eliminate bias. Briefly, cell boundaries were identified by taking the standard deviation of all movement of each cell during acquisition, thresholding and storing the selection area as an ROI. All ROIs were used for measuring intensity of red and green channels for each individual cell and time-point. For cell length measurements, cell ROIs were expanded by two pixels and the region duplicated, and converted to a binary using the Li thresholding algorithm. Individual Feret diameters were measured and recorded for each time-point and ROI. Data was output as .csv files to be further processed (size exclusion < 18 μm or >45 μm), graphed and analyzed. Of the cells that didn’t contract (< 5% contr.) with OT (0.85 nM) alone, approximate 35% did not respond with [Ca^2+^]_i_; this submaximal OT concentration was selected because it produced cell contraction in the majority of cells, without causing complete loss of cell adhesion. Scripts are available at: https://github.com/nickcondon.

### Statistical analysis

Statistical analysis was performed in GraphPad Prism (v7.03). Details of statistical tests used are outlined in each figure legend or supplementary figure legend.

### Data availability

All data are available in the paper. Scripts are freely available on GitHub as stated in Methods.

## Acknowledgments

This work was supported by the National Health and Medical Research Council (1141008 and 1138214), The University of Queensland, the Mater Foundation (Equity Trustees / AE Hingeley Trust) and the National Stem Cell Foundation of Australia. The authors acknowledge the Translational Research Institute (TRI) for the research space, equipment and core facilities that enabled this research. We thank the UQ Biological Resource Facility staff for help with animal work; Dr. Corinne Alberthsen (Mater Research) for assistance with research ethics applications and compliance; Dr. Jerome Boulanger (MRC Laboratory of Molecular Biology) for the 3D denoisingalgorithm; Mr Karsten Bach (University of Cambridge) for assistance with accessing and analyzing RNAseq data; Prof Jane Visvader, Mr Caleb Dawson and Dr Mark Scott for technical assistance with intravital imaging and Mr Eric Pizzani (Translational Research Institute) for research computing support. Samples from the Susan G. Komen Tissue Bank at the IU Simon Cancer Center were used in this study; we thank contributors, including Indiana University (sample collection) as well as donors and their families.

## Supplementary Material

**Fig. S1.**
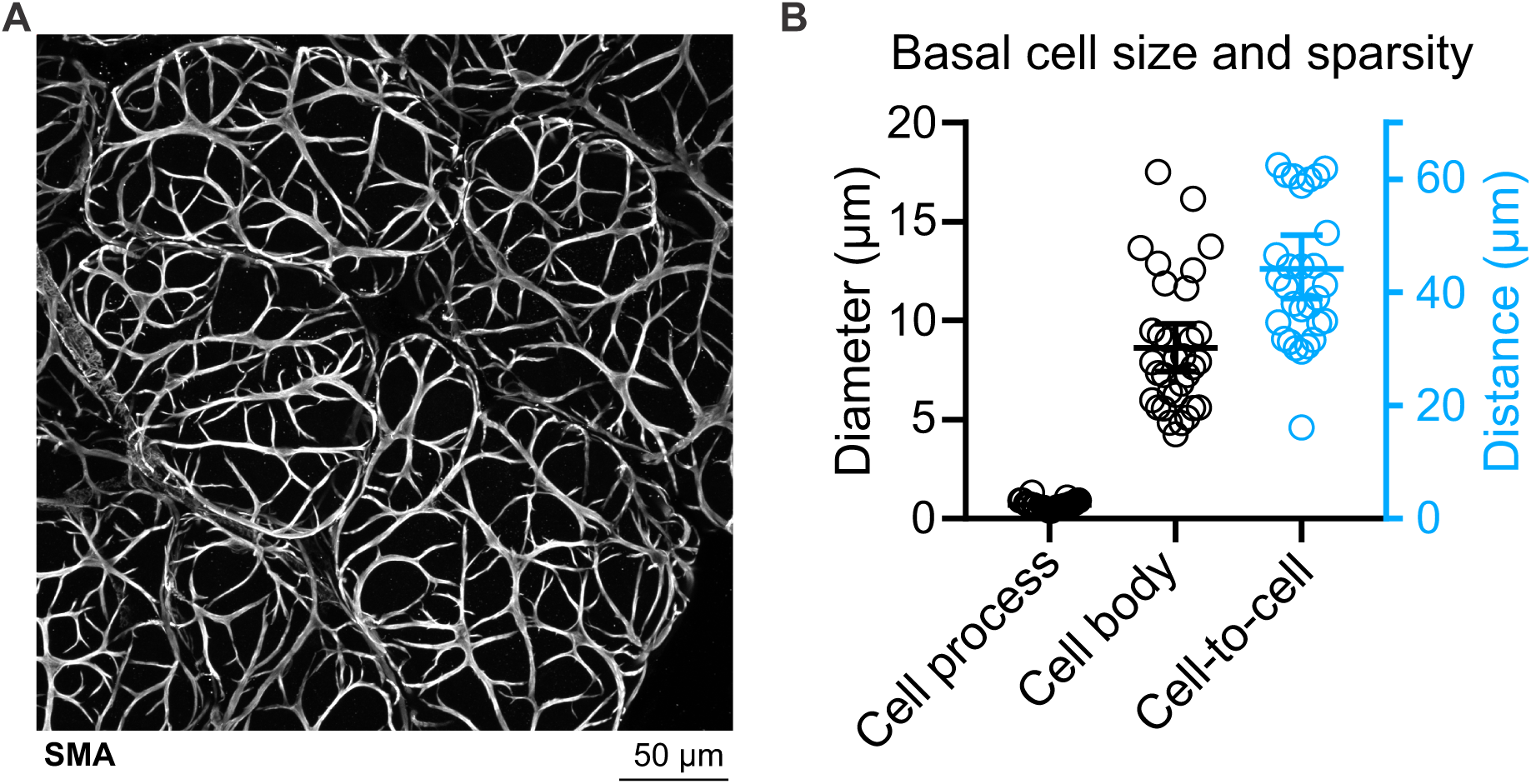
Characterization of morphology and density of mammary alveolar basal cells. (**A**) Maximum intensity z-projection (0-22 μm) of lactating mammary tissue immunostained with smooth muscle actin (SMA) to reveal the cellular morphology and distribution of alveolar basal cells. (**B**) Basal cell diameter at the thinnest cell process and at the cell body (left y-axis). Graph also shows the distance between the body of each basal cell and its closest basal neighbor (right y-axis). Basal cells are separated from their closest neighbor by a distance that is approximately 6× their maximum width. Spatial resolution = 0.3 μm; 33 cells analyzed from n = 3 mice.

**Fig. S2.**
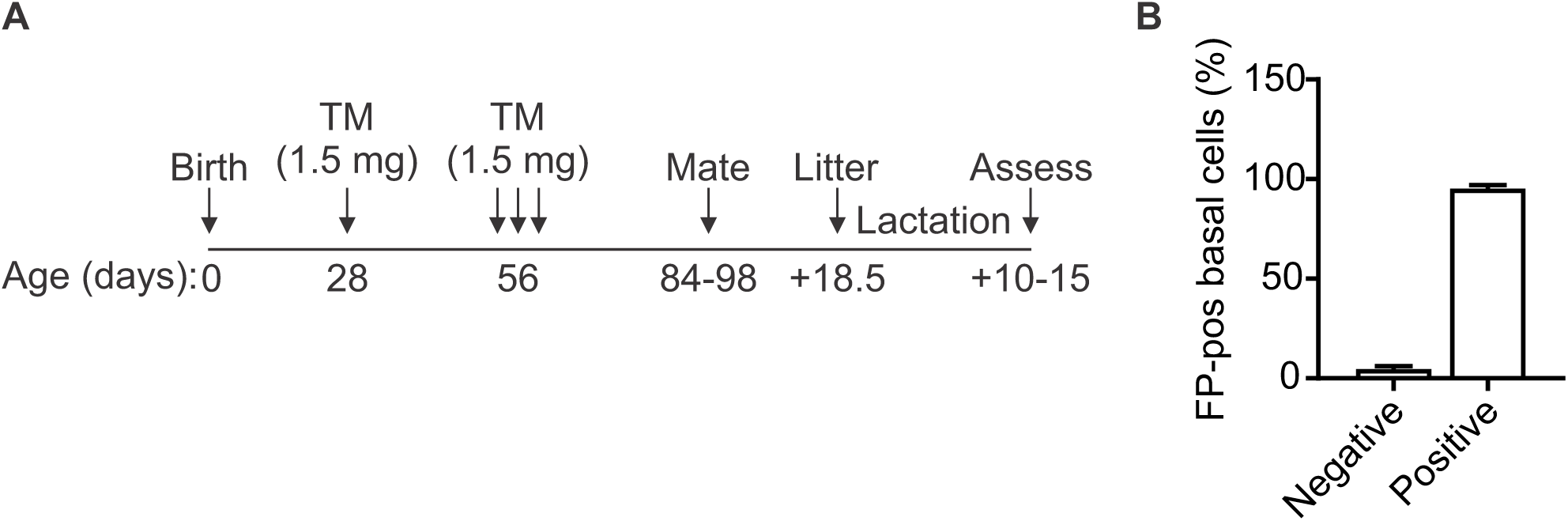
Induction and analysis of fluorescent protein (FP) expression in basal cells. (**A**) Schematic representation of the experimental timeline to generate FP-positive lactating tissue (see Methods). (**B**) The percentage of all basal cells (determined by SMA positivity) that are FP-positive. Positivity was scored from 3D image sequences of cleared tissue; 2400 cells counted from n = 3 mice. TM, tamoxifen.

**Fig. S3.**
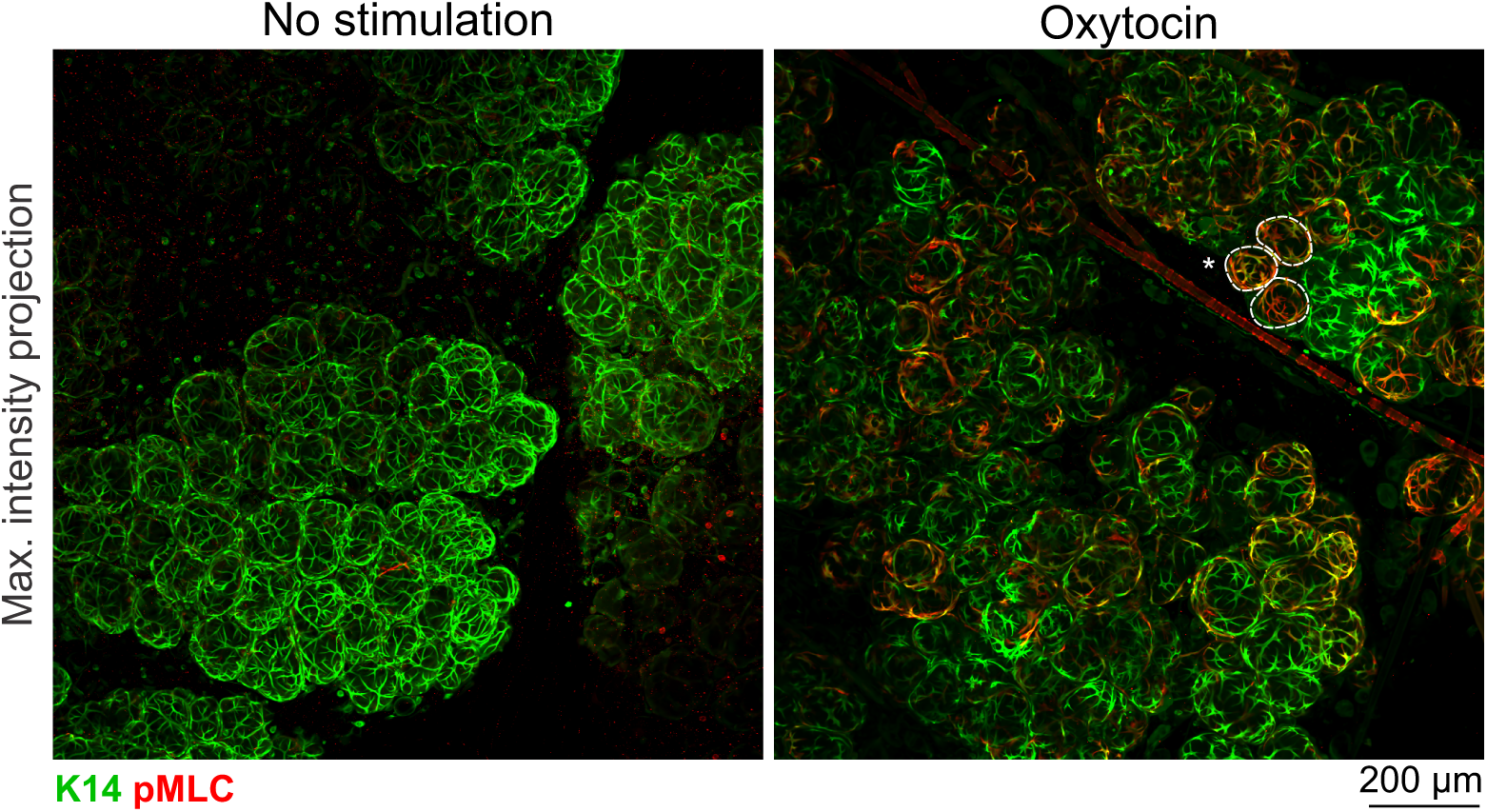
Distribution of pMLC-positive cells. Maximum intensity z-projections [0-80.5 μm (left), 0-140 μm (right)] of cleared mammary tissue immunostained with K14 to reveal basal cells and pMLC to show areas of contractile activity. Dotted line surrounds a cluster of pMLC^+^ cells in lactating mammary tissue stimulated with OT (85 nM) prior to fixation. N = 3.

**Fig. S4.**
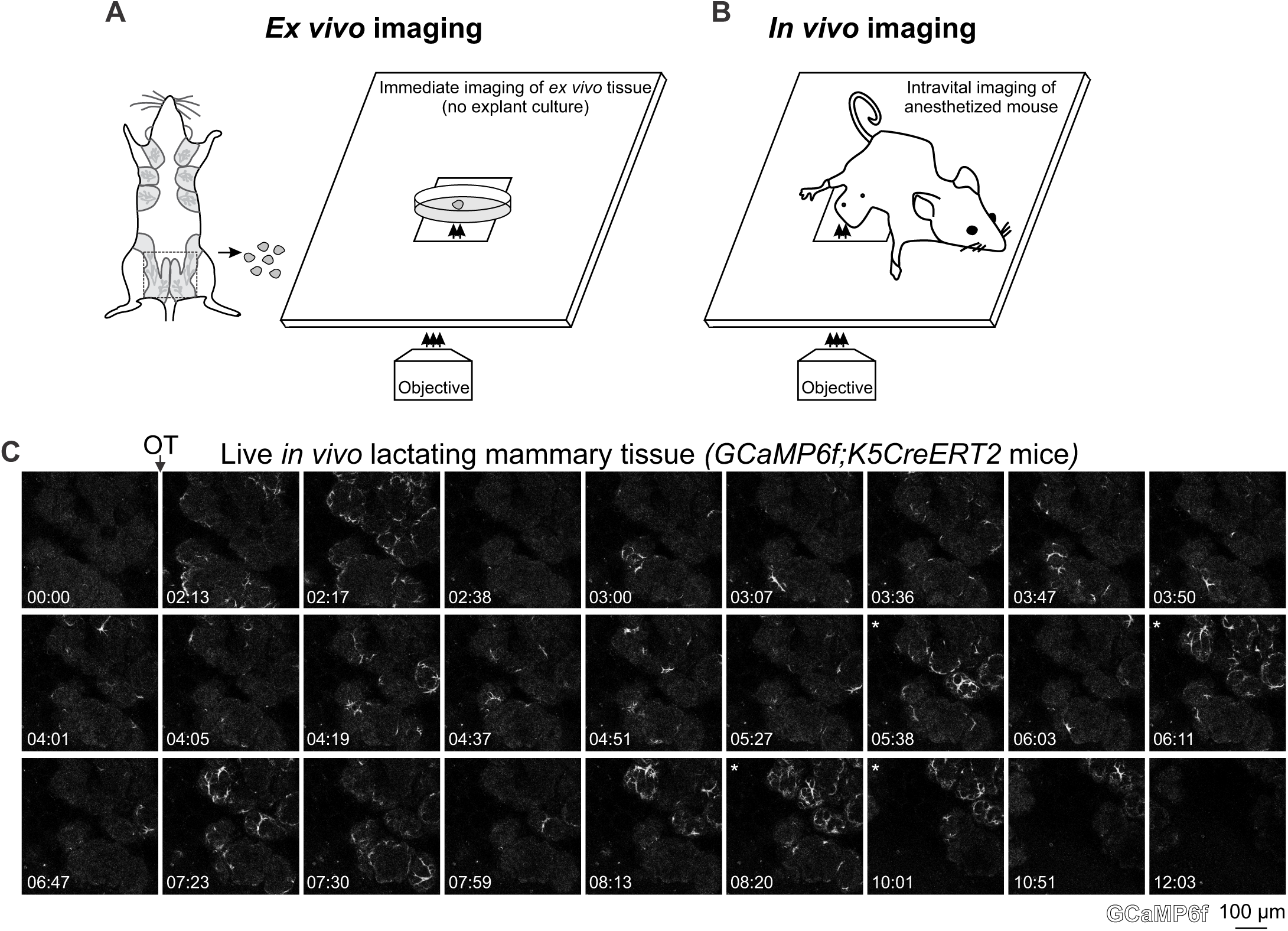
Intravital imaging of OT responses. Imaging set up for *ex vivo* (**A**) and in vivo (**B**) tissue imaging. (**C**) 3D in vivo time-lapse imaging of mammary tissue from a *GCaMP6f;K5CreERT2* lactating mouse injected (i.p.) with OT (2.2 U) between 01:10-01:26 (min:s). Images show maximum intensity *z*-projection 36 μm through tissue. Asterisks show coordinated firing. See also Movie S2.

**Fig. S5.**
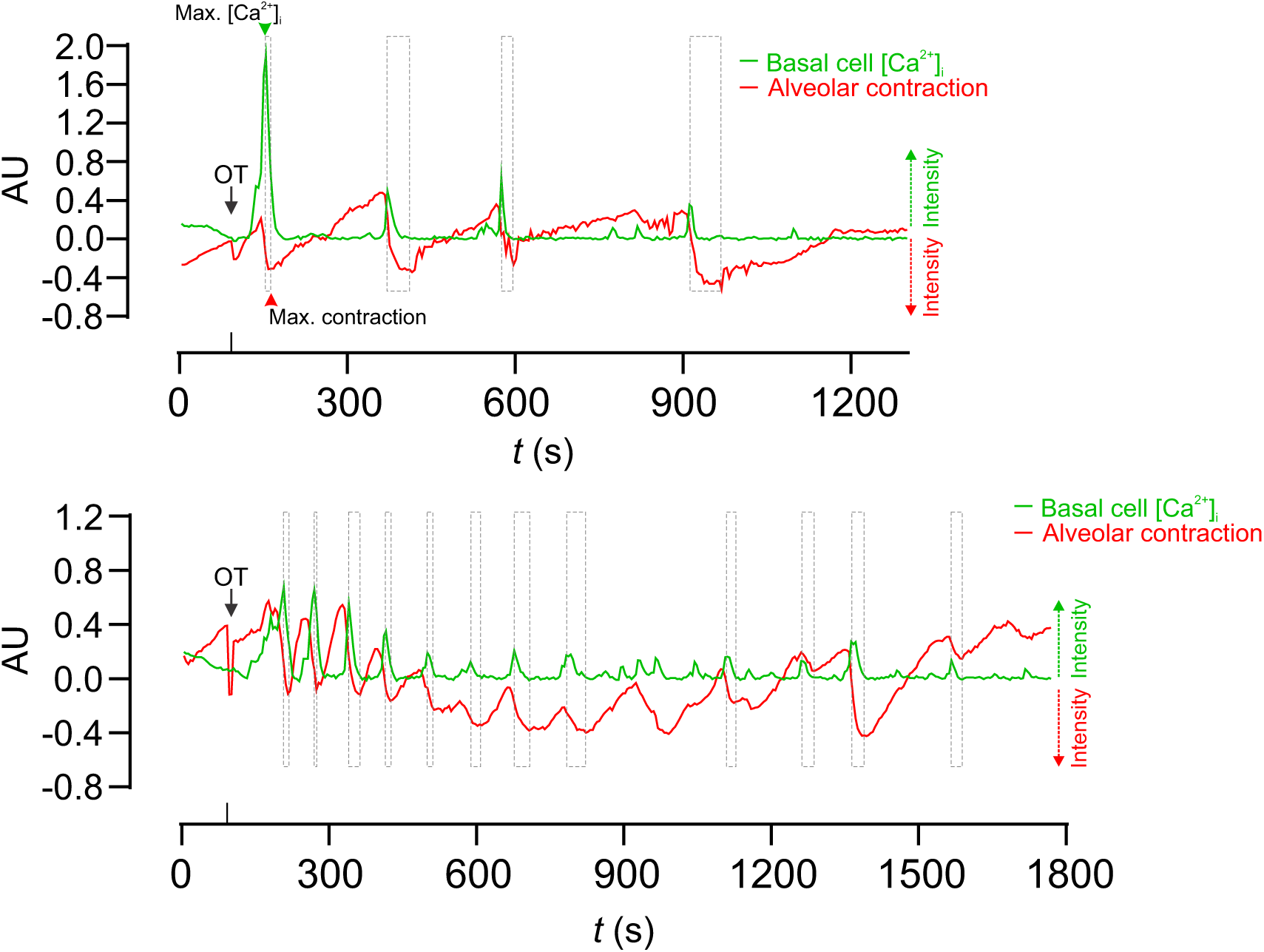
Temporal relationship between [Ca^2+^]_i_ responses and alveolar unit contraction. Additional examples (total n = 3 mice, related to Fig. 1) showing [Ca^2+^]_i_ responses (green) and alveolar unit contractions (red) in lactating mammary tissue from *GCaMP6f;K5CreERT2* mice. [Ca^2+^]_i_ measurements are ΔF/F0. Alveolar unit contractions are shown by negative deflection (reduction in the intensity of the red fluorescence due to displacement of the alveolar unit). Boxes align with the peak [Ca^2+^]_i_ response (left) and the peak contractile response (right). AU, arbitrary unit.

**Fig. S6.**
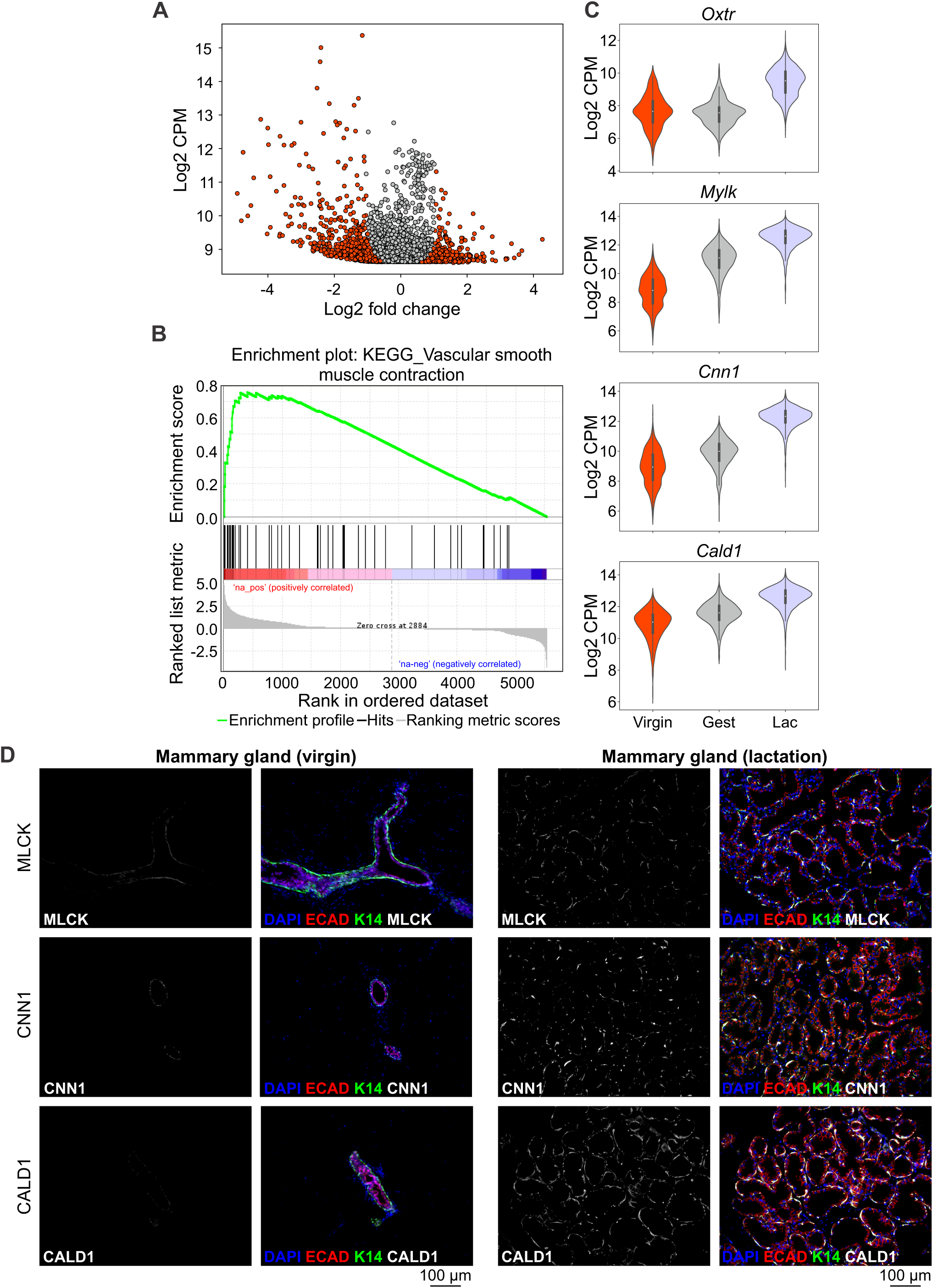
Upregulation of the contraction toolkit in basal cells during pregnancy and lactation. (**A**) Volcano plot showing differential expression (red dots; FDR < 0.05) from single cell RNA sequencing data [Basal-Virgin vs. Basal-Lac (lactation) clusters; Bach et al., (2017) Nat Commun]. (**B**) Genes in the KEGG vascular smooth muscle contraction pathway showed significant enrichment in Basal-Lac vs. Basal-Virgin by GSEA using MSigDB curated gene sets. (**C**) Frequency distribution of expression values from Basal-Virgin, Basal-Gest (gestation) and Basal-Lac clusters shown as a violin plot for each gene. Expression (y-axis) shown as log2 counts per million (CPM); width reflects the distribution of values on the y-axis. (**D**) Immunohistochemical staining for myosin light chain kinase (MLCK), calponin (CNN1) and caldesmon (CALD1) in mammary tissue isolated from virgin and lactating wild-type mice. E-cadherin (red) shows the luminal cell lineage; K14 (green) shows the basal cell lineage. Nuclei are stained with DAPI (blue); n = 3 mice.

**Fig. S7.**
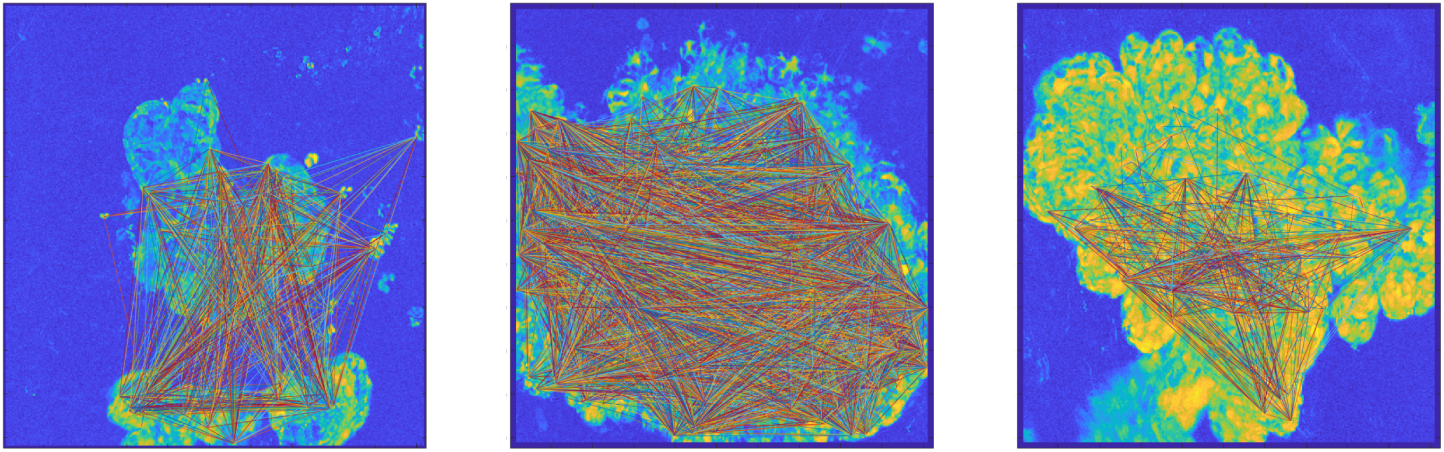
Tissue coordination in mammary tissue from pregnant (15.5-16.5 d.p.c) *GCaMP6f-TdTom;K5CreERT2* mice. Graph theory showing connection between highly correlated events; n = 3 mice (gestation).

**Fig. S8.**
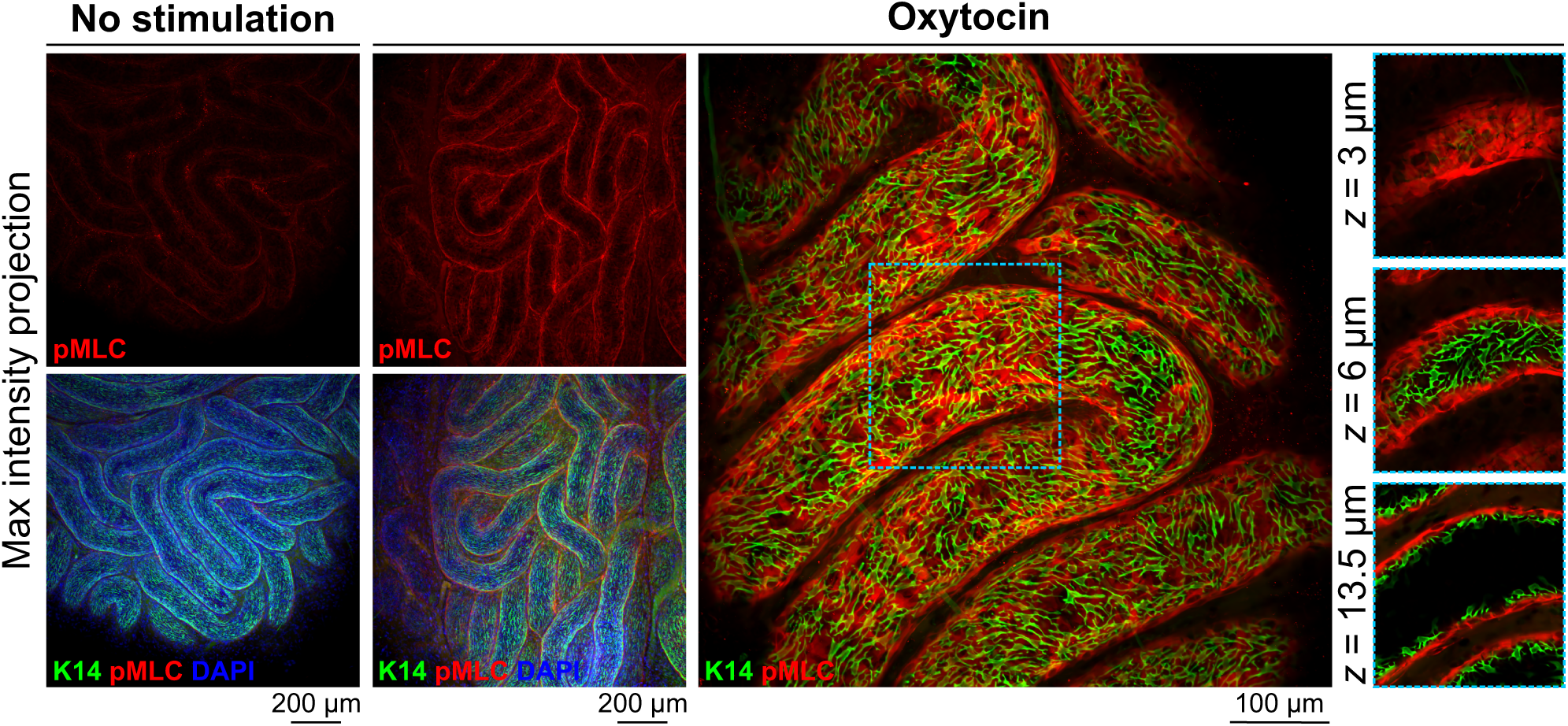
Detection of activated myosin in epididymal tissue. Maximum intensity z-projections of cleared epididymal tissue immunostained with K14 (green) and pMLC (red). Tissue was stimulated with OT (850 nM) 20 min prior to fixation, as indicated. Nuclei (DAPI) are blue; n = 3 mice.

**Fig. S9.**
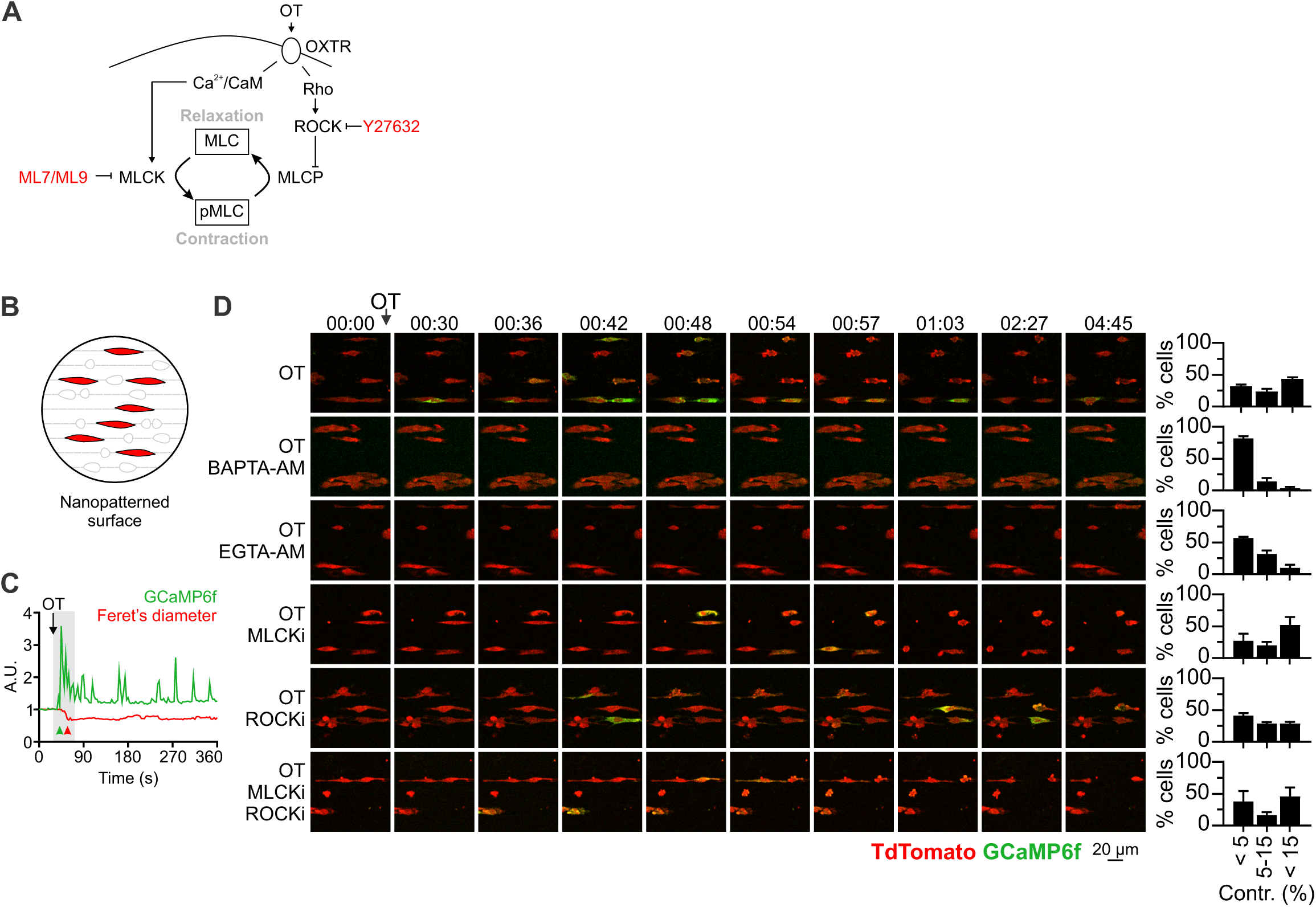
Pharmacological inhibitors of contractile machinery. (**A**) Schematic representation of contractile pathways in smooth muscle cells and (**B**) the assay developed to assess in vitro contractile responses. (**C**) First phase (gray box) Ca^2+^-contraction coupling in primary cells in vitro. (**D**) 2D time-lapse imaging of GCaMP6f-TdTomato dual positive cells in response to OT (0.85 nM, 00:27), or with OT in cells loaded with the intracellular Ca^2+^ chelators (BAPTA-AM and EGTA-AM), MLCKi (ML-7), ROCKi (Y27632) or a combination of MLCKi and ROCKi. Graphs show average (± SEM) percent cells in each bin [<5%, 5-15%, >15% reduction in Feret’s diameter (contr.)]; 80-150 cells analyzed for each treatment from n = 3 mice.

**Fig. S10.**
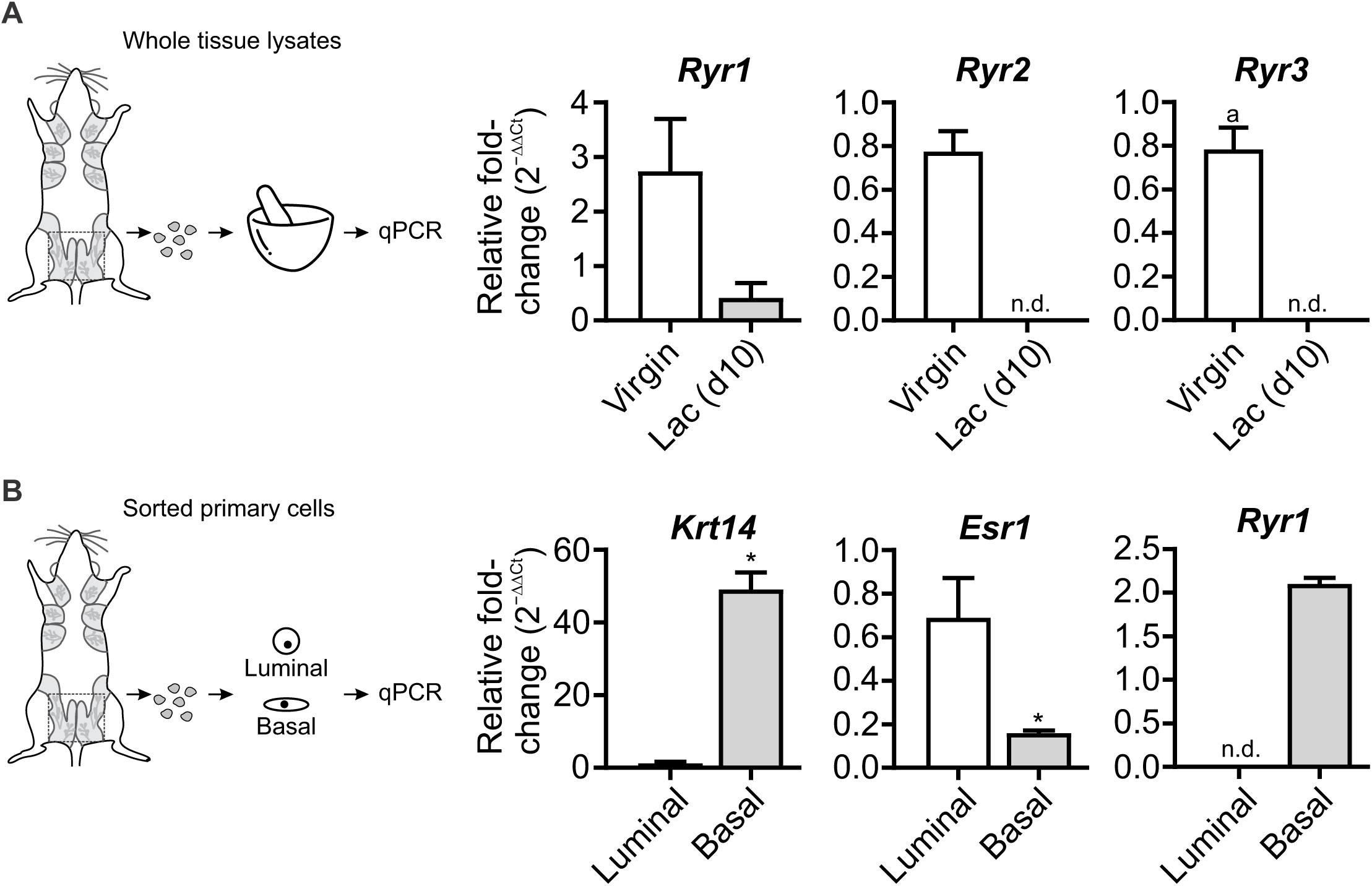
mRNA levels of ryanodine receptors. (**A**) *Ryr1*, *Ryr2* and *Ryr3* expression in lysates prepared from whole mammary tissue (including luminal, basal and stromal cells) dissected from virgin or lactating animals (n = 4 mice). (**B**) *Krt14*, *Esr1* and *Ryr1* levels in freshly sorted luminal and basal cells (n = 3 mice). Graphs show mean ± SEM; * P < 0.05 (Student’s t-test); n.d., not detected. *Ryr2* and *Ryr3* transcripts were either not detected or detected at very low levels in only a fraction of samples from both luminal and basal cells.

**Fig. S11.**
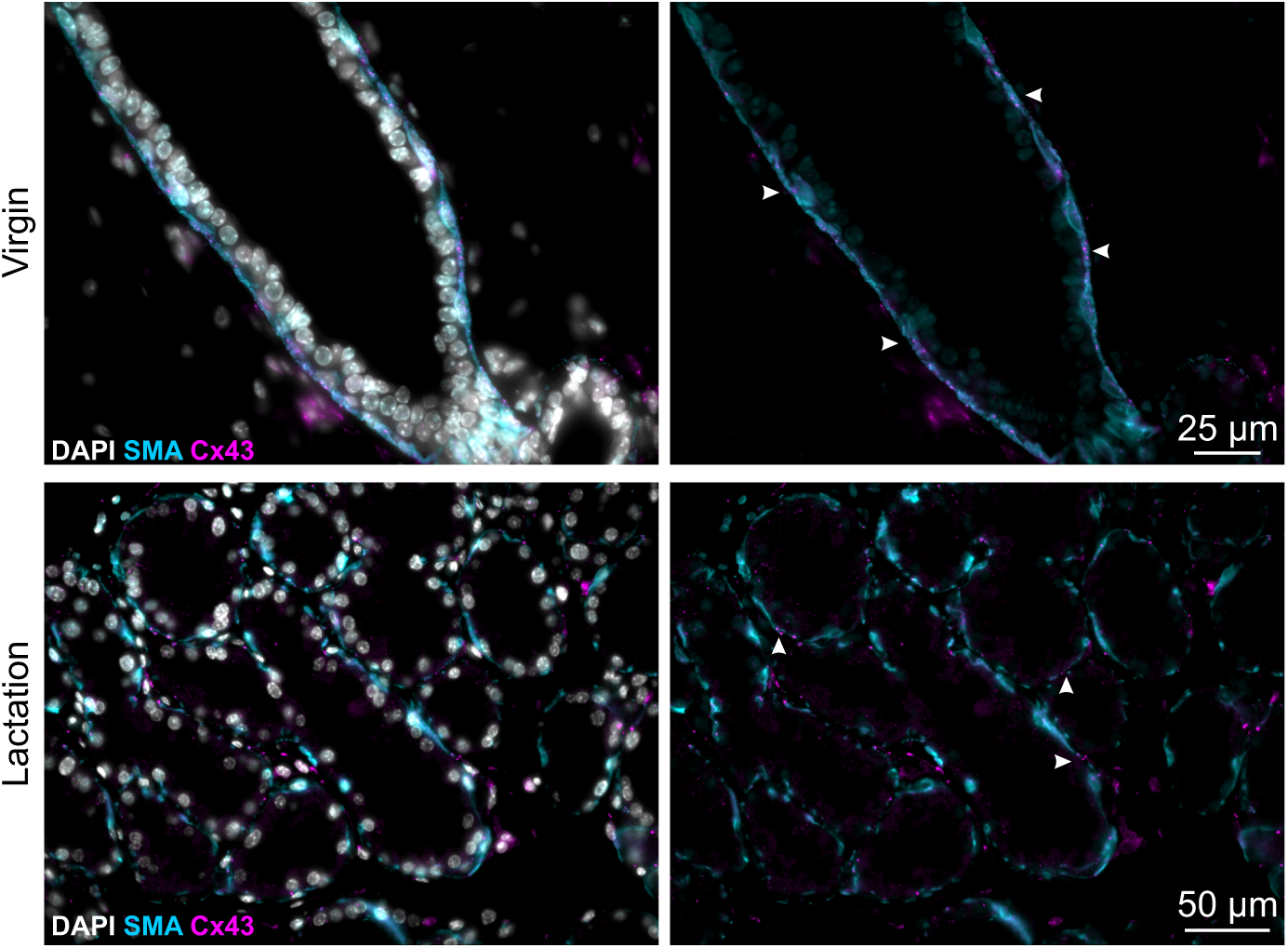
Cx43 expression (2D tissue sections). Cx43 immunostaining (magenta) in virgin and lactating mammary tissue. SMA (cyan) shows the basal cell lineage. Nuclei are stained with DAPI (gray).

## Supplementary Movie

Movie S1.

OT response in live mammary tissue isolated from a lactating *GCaMP6f;K5CreERT2* mouse. GCaMP6f (green) and CellTracker™ (red). OT was added at 01:33 (min:s). Total movie length is 29:30 (min:s). Related to Fig. 1.

Movie S2.

Intravital imaging of OT response in a lactating *GCaMP6f;K5CreERT2* mouse. OT was injected between 01:10-01:26 (min:s) and exhibits a similar (albeit faster) response to that seen acute ex vivo tissue (**Movie S1**). Total movie length is 29:30 (min:s). Related to Fig. S4.

Movie S3.

OT response in live mammary tissue isolated from a lactating *GCaMP6f-TdTom;K5CreERT2* mouse. GCaMP6f (green) and TdTomato (red). OT was added at 01:09 (min:s). Total movie length is 23:03 (min:s). Related to Fig. 2.

Movie S4.

OT response in live mammary (alveolar) tissue isolated from a pregnant (15.5-16.5 d.p.c.) *GCaMP6f-TdTom;K5CreERT2* mouse. GCaMP6f (green) and TdTomato (red). OT was added at 00:38 (min:s). Total movie length is 25:08 (min:s).

Movie S5.

OT response in live mammary (alveolar) tissue isolated from a pregnant (15.5-16.5 d.p.c.) *GCaMP6f-TdTom;K5CreERT2* mouse. Tissue was incubated in Ca^2+^ free buffer prior to imaging. Extracellular Ca^2+^ was added back at a final concentration of 1 mM (free Ca^2+^), as indicated. Total movie length is 29:55 (min:s). Related to Fig 2.

Movie S6.

OT response in live mammary (ductal) tissue isolated from a pregnant (15.5-16.5 d.p.c.) *GCaMP6f-TdTom;K5CreERT2* mouse. GCaMP6f (green) and TdTomato (red). OT was added at 01:15 (min:s), as indicated. Total movie length is 26:13 (min:s). Related to Fig. 3.

Movie S7.

OT response in live mammary (ductal) tissue isolated from a lactating *GCaMP6f-TdTom;K5CreERT2* mouse. GCaMP6f (green) and TdTomato (red). OT was added approximately 00:45 (min:s) prior to imaging. Total movie length is 26:13 (min:s). Related to Fig. 3.

Movie S8.

OT response in live lacrimal tissue isolated from a *GCaMP6f-TdTom;K5CreERT2* mouse. GCaMP6f (green) and TdTomato (red). OT was added at 00:45 (min:s), as indicated. Total movie length is 15:55 (min:s). Related to Fig. 4.

Movie S9.

OT response in live epididymal tissue isolated from a *GCaMP6f-TdTom;K5CreERT2* mouse. GCaMP6f (green) and TdTomato (red). OT was added at 01:38 (min:s), as indicated. Total movie length is 25:48 (min:s). Related to Fig. 4.

Movie S10.

Tissue contraction in response to pharmacological inhibitors. From top to bottom: uterus, epididymis, bladder and mammary gland loaded with CellTracker^TM^. From Left to right: control, MLCKi+ROCKi, MLCKi+ROCKi+PKCi+CaMKIIi. Uterus and mammary tissue was stimulated with OT (85 nM). Epididymal tissue was stimulated with OT (850 nM). Bladder tissue was stimulated with carbachol (10 μM) at the commencement of imaging. Related to Fig. 5.

Movie S11.

OT response in live primary cells isolated from a pregnant (15.5-16.5 d.p.c.) *GCaMP6f-TdTom;K5CreERT2* mouse and grown on a nanopatterned surface. OT was added at 00:27 (min:s), as indicated. Total movie length is 06:00 (min:s). Related to Fig. S9.

Movie S12.

OT response in live primary cells isolated from a pregnant (15.5-16.5 d.p.c.) *GCaMP6f-TdTom;K5CreERT2* mouse, grown on a nanopatterned surface and pre-treated with BAPTA-AM for 30 min. OT was added at 00:27 (min:s), as indicated. Total movie length is 06:00 (min:s). Related to Fig. S9.

Movie S13.

OT response in live mammary tissue isolated from a lactating *GCaMP6f;K5CreERT2* mouse. Tissue was pre-treated with dantrolene (20 μM) for approx. 5 h. GCaMP6f (green) and TdTomato (red). OT was added at 01:53 (min:s). Total movie length is 32:51 (min:s). Related to Fig. 6.

Movie S14.

OT response in live mammary tissue isolated from a lactating *GCaMP6f;K5CreERT2* mouse. Tissue was pre-treated with ryanodine (100 μM) for approx. 5 h. GCaMP6f (green) and TdTomato (red). OT was added approx. 12 min prior to imaging. Total movie length is 12:14 (min:s).

Movie S15.

The effect of nifedipine on tissue synchronization promoted by dantrolene. Tissue from a pregnant *GCaMP6f-TdTom;K5CreERT2* mouse was pre-treated with dantrolene and stimulated with OT (85 nM) approx. 12 min prior to imaging. Nifedipine (20 μM) was added at 21:30 (min:s), as indicated. Total movie length is 29:55 (min:s). Related to Fig. 6.

